# Deep analysis of the Major Histocompatibility Complex associations using covariate analysis and haploblocks unravels new mechanisms for the molecular etiology of Elite Control in AIDS

**DOI:** 10.1101/2024.12.07.625684

**Authors:** Myriam Rahmouni, Sigrid Le Clerc, Jean-Louis Spadoni, Taoufik Labib, Maxime Tison, Raissa Medina-Santos, Armand Bensussan, Ryad Tamouza, Jean-François Deleuze, Jean-François Zagury

## Abstract

**Introduction:** We have reanalyzed the genomic data from the International Collaboration for the Genomics of HIV (ICGH), focusing on HIV-1 Elite Controllers (EC).

**Methods:** A genome-wide association study (GWAS) was performed, comparing 543 HIV-1 EC individuals with 3,272 uninfected controls (CTR) of European ancestry. 8 million single nucleotide polymorphisms (SNPs) and HLA class I and class II gene alleles were imputed to compare EC and CTR.

**Results:** 2,626 SNPs were associated with EC (p<5.10-8), all located within the Major Histocompatibility Complex (MHC) region. Stepwise regression analysis narrowed this list to 17 SNPs. In parallel, 22 HLA class I and II alleles were associated with EC. Through meticulous mapping of the LD between all identified signals and employing reciprocal covariate analyses, we delineated a final set of 6 independent SNPs and 3 HLA class I gene alleles that accounted for most of the associations observed with EC. Our study revealed the presence of cumulative haploblock effects (SNP rs9264942 contributing to the HLA-B*57:01 effect) and that several HLA allele associations were in fact caused by SNPs in linkage disequilibrium (LD). Upon investigating SNPs in LD with the selected 6 SNPs and 3 HLA class I alleles for their impact on protein function (either damaging or differential expression), we identified several compelling mechanisms potentially explaining EC among which: a multi-action mechanism of HLA-B*57:01 involving MICA mutations and MICB differential expression overcoming the HIV-1 blockade of NK cell response, and overexpression of ZBTB12 with a possible anti-HIV-1 effect through HERV-K interference; a deleterious mutation in PPP1R18 favoring viral budding associated with rs1233396.

**Conclusion:** Our results show that MHC influence on EC likely extends beyond traditional HLA class I or class II allele associations, encompassing other MHC SNPs with various biological impacts. They point to the key role of NK cells in preventing HIV-1 infection. Our analysis shows that HLA-B*57:01 is indeed associated with a partially functional NK cell response which could also explain this marker’s involvement in other diseases such as psoriasis. More broadly, our findings suggest that within any HLA class I and II association in diseases, there may exist distinct causal SNPs within this crucial, gene-rich, and LD-rich MHC region.

## Introduction

Despite the availability of effective antiretroviral drugs, HIV-1 infection continues to be an important public health concern, with a large number of new infections and deaths annually, particularly in low-income countries (1). In the early 2000s, a subset of HIV-positive individuals who maintained consistently low viral loads for multiple years without receiving any treatment were recognized as Elite Controllers (EC). These individuals represent approximately 0.2 to 0.5% of the HIV-positive population (2), including in African cohorts (3), a finding that has been confirmed in the GRIV cohort comprising long-term non progressors and EC (4,5). Previous genome-wide association studies (GWAS) have shown that genetic variants within the Major Histocompatibility Complex (MHC) region exert the most significant influence on viral load control and disease progression (4,6,7), and particularly for the Elite Control phenotype. Given that EC subjects possess natural control over viral infection, they represent a valuable population for investigating the molecular mechanisms of protection. So far, the explanations for the biological impact of the MHC region in HIV-1 control have been limited to the presentation role of the classical HLA class I alleles, HLA-A, HLA-B, and HLA-C (8–11). In 2010, Pereyra et al. published a study on 2000 viral controllers (7) and identified 323 signals associated with the control of viral load, and more specifically 4 SNP by stepwise regression, that seemed to impact independently the control of viral load in the MHC region. In a recent study, we have conducted an extensive GWAS comparing 543 individuals characterized as HIV Elite Controllers (ECs) with 3272 uninfected control subjects of European descent (12). This group of 543 elite controllers is very powerful since it corresponds to the extreme phenotype of a cohort larger than 100,000 infected individuals. Leveraging on the latest bioinformatics databases, we have imputed 8M SNPs over the genome and identified 2,626 significant signals surpassing the genome-wide significance threshold (5.10^-8^), all in the MHC region. It is noteworthy that our case-control study has identified 8 times more signals than the 323 signals previously identified by Pereyra et al. and this is likely due to the more extreme phenotype analyzed (elite controllers instead of viral controllers) and possibly in part to the progress of the SNP database since 14 years. This very large number of associations witnesses the extensive LD that characterizes the MHC locus. The MHC region spans approximately 5 million base pairs (genomic coordinates 28,477,797 to 33,448,354) on chromosome 6, according to the latest GENCODE gene annotation for the GRCh38 reference genome (13). Within this region, a total of 373 protein-coding genes, 18 pseudogenes, and 12 non-coding RNA genes were identified by the GENCODE annotation. The human MHC houses several genes crucial for both innate and adaptive immune responses. Variations in Human Leukocyte Antigen (HLA) class I alleles have been associated with HIV elite control in both European and African populations, and notably the HLA-B*57 allele has been associated with an enhanced control over HIV-1 infection. In our recent study, we had focused our analysis on the HLA-B*57 class I allele and described for the first time a very large haploblock corresponding to the HLA-B*57:01 allele, spanning 1.9 MB (12). We have shown how this haploblock could impact HIV-1 replication through the existence of mutations or through the differential expression of numerous proteins (12).

In the present study, following the GWAS comparing EC and CTR individuals, we have performed a stepwise regression analysis in order to narrow down the 2626 identified signals to a smaller number of independent SNPs. Our goal was also to compare the resulting SNPs with the classical HLA class I and class II alleles, and to look for their relative impact on the EC phenotype.

## Results

### GWAS and stepwise regression

The 2626 signals of the GWAS comparing EC with CTR are illustrated in Figure 1, they are all located in the MHC region. Similar to the approach taken by Pereyra et al. (7), we have conducted a stepwise regression in our case-control study and identified 17 SNPs significantly associated with Elite Control. These 17 SNPs, the p-values and the odds ratios (OR) from the original GWAS as well as from the stepwise regression are detailed in Table 1. The 17 SNP localization is shown in supp Figure 1 and they cover a large portion of the MHC region. When looking at Table 1, the 17 SNPs demonstrate diverse trajectories: rs1233396 maintains an OR < 1 in both the original GWAS and the stepwise analysis; rs9262549 transitions from an OR < 1 in the original GWAS to an OR > 1 in the stepwise analysis; three SNPs (rs2596485, rs13208886, rs28732175) shift their OR from > 1 to < 1 in the stepwise analysis; and two SNPs (rs112630608 and rs111301312) exhibit a reduction in OR by a factor of 2 compared to the OR in the original GWAS, with the remaining SNPs showing minimal changes. Notably, rs111301312, which tags the well-known HLA-B*57:01 allele, displays a marked decrease in the stepwise analysis.

**Figure 1.**
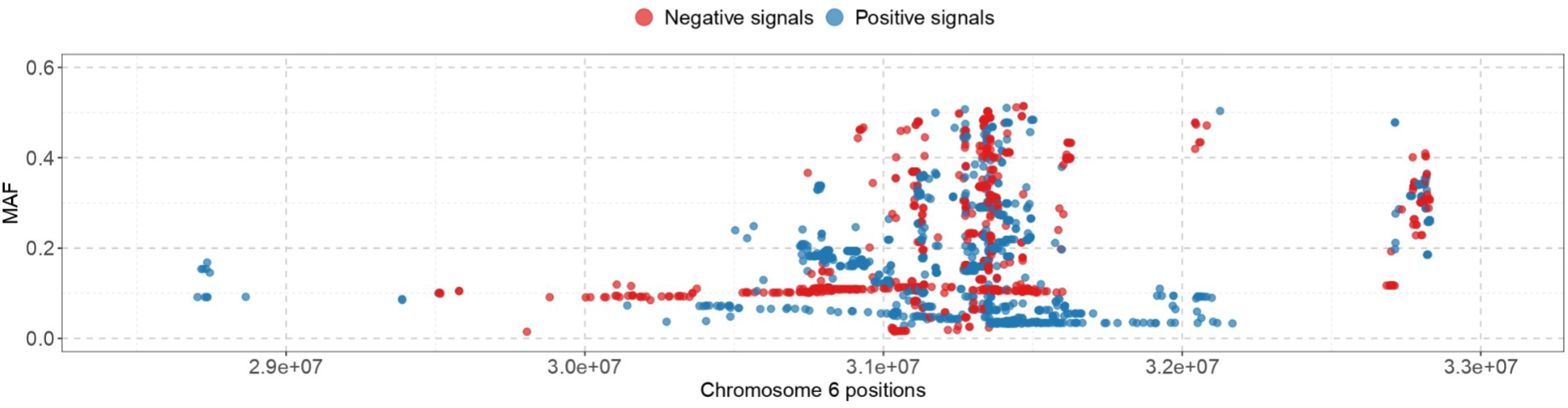
Representation of the 2626 SNPs associated with EC in the MHC region with their localization in chromosome 6 GRCh38) (x-axis) and their MAF (y-axis) Representation of the 2626 SNPs significantly associated (p<5. 10^-8^) with elite control in the EC vs CTR GWAS. In blue, the SNPs whose minor allele favors EC. In red, the SNPs whose minor allele prevents EC.

**Table 1.**
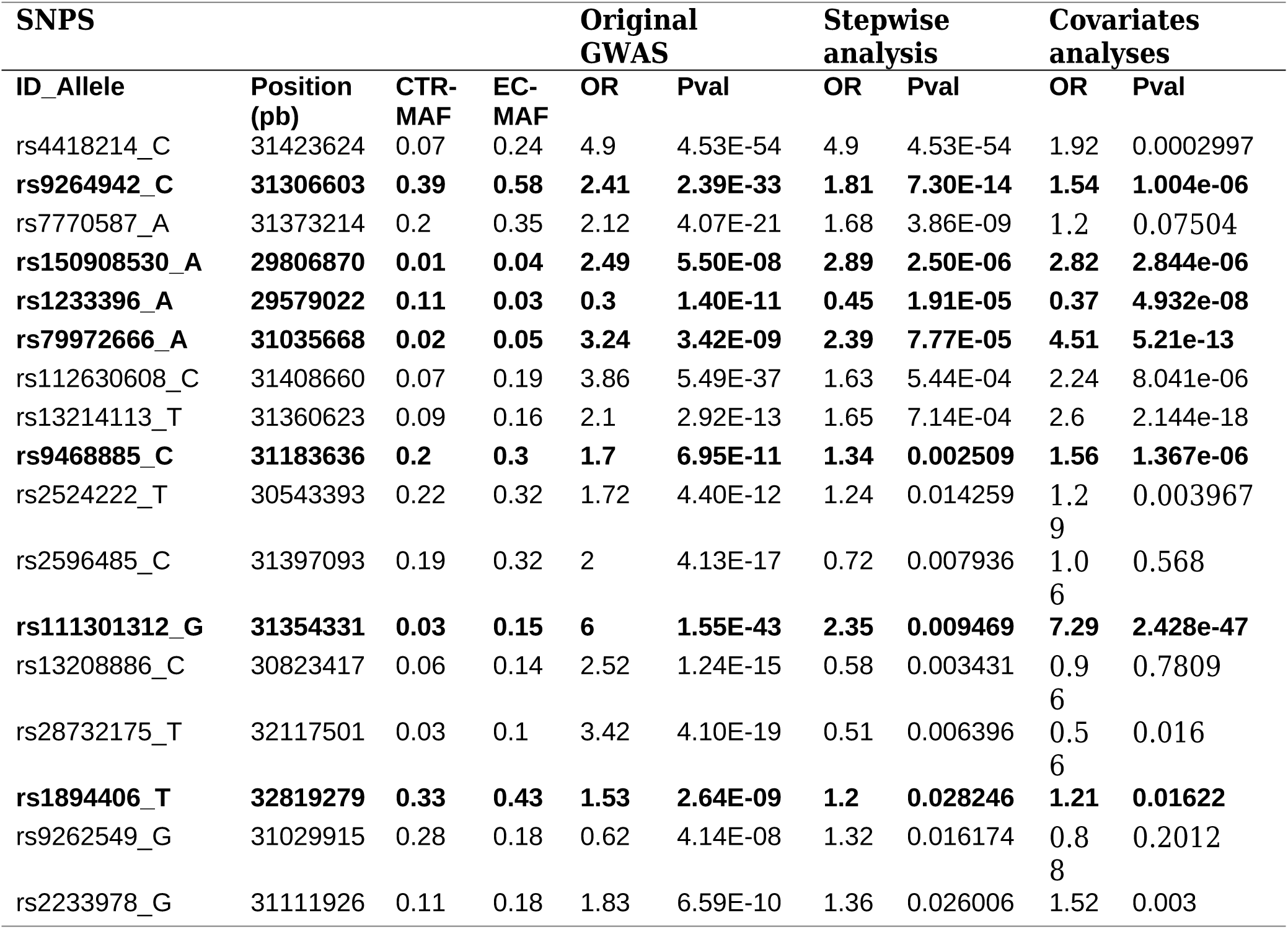
SNPs remaining after stepwise regression and covariate analysis. Table giving the MAF, OR, and p values of the SNPs obtained by stepwise regression analysis. The OR and p values from the original GWAS are also provided. The covariate analysis made with the leaves of the tree (Figure 2A) has eliminated 4 SNPs. The covariate analysis made with the significant HLA alleles has eliminated 6 additional SNPs. The 7 selected SNPs remaining after all covariate analyses are in bold.

### LD Links between the SNPs identified by stepwise regression

The investigation of LD among these 17 single nucleotide SNPs within the control population (CTR) reveals a substantial degree of LD (see Supplementary Table 1 and Figure 2A). The tree representing the SNP minor allele links within the CTR population illustrates that each terminal node (leaf) of the tree, that corresponds to the SNPs with the smallest minor allele frequency (MAF), may influence the association study either independently or through SNPs located in the branches associated with this leaf, effectively through their haploblock. As an illustration, SNP rs9264942, which exhibits the highest MAF, occupies a central position in the tree due to its strong LD with the majority of the other SNPs (Figure 2A and Supplementary Table 1). Interestingly, its minor allele encompasses the minor allele of rs111301312 – tag of HLA-B*57:01-(D’=1). That possesses the highest OR in the original GWAS (OR=6, see Table 1). There are 2 possibilities: either the minor allele of rs9264942 exerts a statistical effect on elite control independently, or its OR is attributable to the cumulative effects of the ORs of the minor alleles it encompasses. To investigate this, we performed an initial analysis by calculating the association values (OR and p-value) for all 17 SNPs while using the SNPs that represent the leaves of the tree as covariates in the regression analysis (highlighted in blue in Figure 2A). The rationale for selecting these leaves is their lack of LD with one another, thereby facilitating the regression analysis without confounding effects. For rs9264942, we observed an association with an OR of 1.54, slightly smaller than that identified in the original GWAS regression but still significant (see Table 1, column on covariate analyses). Remarkably, when incorporating the 16 other SNPs as covariates in the regression, a similar OR of 1.56 was obtained (see Table 1). This finding suggests that the association identified for rs9264942 is independent of the other SNPs represented in the tree. Consequently, the strongest effect observed for rs111301312 (OR=6) is evidently a result of the combined influence of rs9264942 (OR=1.56) and one or more SNPs located within its branch (i.e., within its haploblock), as the OR derived from stepwise regression for rs111301312 is 2.35 (Table 1, column on stepwise regression). In a second example, the only two SNPs associated with a negative effect on elite control in the original GWAS (rs1233396 and rs9262549, both with OR<1) are in LD with one another (Figure 2A and Supplementary Table 1). In a third example, one of the 3 SNPs exhibiting positive effects in the original GWAS that subsequently show negative effects in the stepwise regression (rs13208886) is in high LD with the strongest signal rs111301312 (Figure 2A and Supplementary Table 1). This explains its protective effect in the original GWAS. However, when rs111301312 is included as a covariate, its protective effect disappears (Table 1).

**Figure 2.**
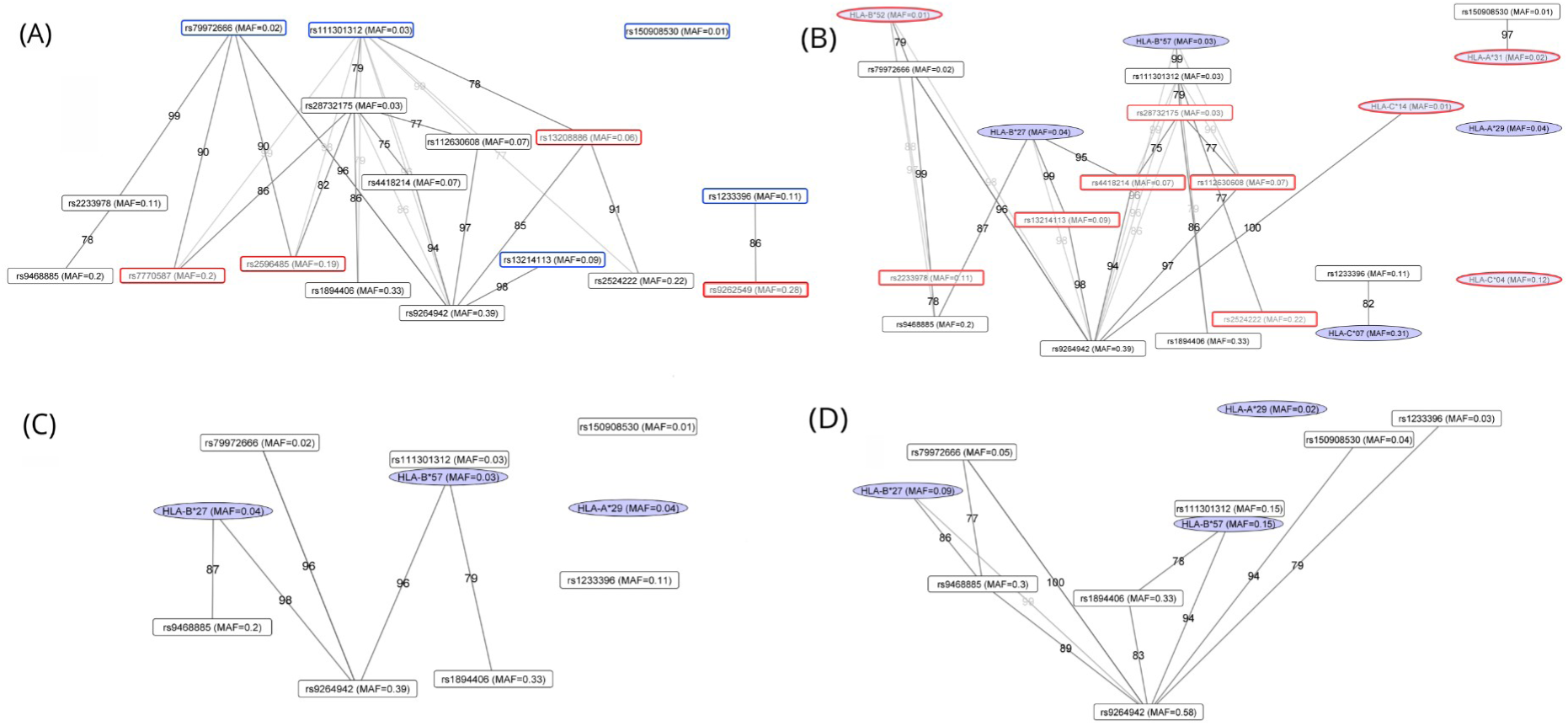
Representation of the stepwise regression SNP and HLA alleles associated with EC. The SNP/HLA alleles are positioned in the tree according to their MAF from higher MAF (bottom) to lower MAF (top). The links between the SNPs are marked by lines and by numbers indicating the percentage of individuals containing the minor alleles of the upper SNP also containing the minor alleles of the lower SNP/HLA allele. This LD links have been computed from the CTR population. **A.** Tree representation of the 17 SNPs alone. The SNPs framed in blue are the leaves used to eliminate redondant SNPs by covariate analysis. The SNPs framed in red are the ones eliminated after the first round of covariate analysis. (Table 1). **B.** Tree representation of the 13 SNPs remaining after the first covariate analysis, and the 8 significant HLA alleles remaining after the analysis of LD between the 22 associated HLA alleles (Table 2). These SNP and HLA alleles have been analyzed by covariate analysis and the ones framed in red are the ones eliminated because dependent on the others (Supplementary Table 3). **C.** Tree representation of the 9 SNP and HLA alleles remaining after covariate analysis, that exert independent statistical effects on elite control. The numbers correspond to the percentage linking the SNP minor alleles or HLA alleles, and have been computed from the CTR population. **D.** Tree representation of the 9 SNP and HLA alleles selected in Figure 3C, based on their links computed directly from the EC population.

Using this approach of assessing the impact of the 5 SNP leaves as covariates, we were able to keep 13 independent SNPs with a significant signal in the regression analysis (Table 1, column covariate analyses), while excluding 4 others. The revised tree composed of these 13 SNPs is shown in Figure 2B. Among these 13 SNPs, we find rs1233396 preventing Elite Control in the original GWAS, our analysis showing that this association was not a simple mirror effect of the rs111301312/HLA-B*57:01 haploblock.

### HLA class I and class II allele imputation and associations with EC

We imputed the HLA class I and class II alleles using the software SNP2HLA (14). We then performed the comparison of EC with CTR for the HLA alleles. Table 2A displays the 13 protective associations of HLA class I and class II alleles, consistent with prior studies comparing EC cohorts to controls (10). Notable signals include HLA-B*57 (p=1.7 10-49), HLA-C*06 (p=2.2 10-24), HLA-B*27 (p=1.3 10-15), HLA-A*31 (p=1.5 10-6), and HLA-B*52 (p=2.5 10-6). Likewise, Table 2B outlines the 9 HLA class I and II alleles having a significant preventive impact on EC, with strong signals such as HLA-C*07 (p=1.1 10-19) and HLA-B*08 (p=7.1 10-10). As previously done with the SNPs, we assessed the LD between the 22 HLA class I and class II signals of Tables 2A and 2B in the CTR population (Supplementary Table 2) and performed a regression analysis incorporating various HLA alleles as covariates (Supplementary Table 3). For instance, the signal of HLA-C*06 disappeared upon including HLA-B*57:01 as a covariate (Supplementary Table 3), in line with the fact that HLA-B*57:01 and HLA-C*06 are part of an ancestral haplotype (15–17). Similarly, the signal for HLA-B*08 disappeared when HLA-C*07 was included as a covariate (Supplementary Table 3), suggesting a haplotypic relationship, as HLA-B*08 is significantly less prevalent while always associated with HLA-C*07. Following this covariate analysis, several HLA alleles could be excluded (Supplementary Table 3), leading to the identification of 8 independent HLA class I signals, comprising 5 protective and 3 preventive associations with EC (highlighted in Table 2).

**Table 2.** HLA class I and class II alleles associated with EC. Table providing MAF, OR and p values for the 22 HLA class1 and class II alleles associated with EC. Some of the alleles have been eliminated by various steps of covariate analysis (Supplementary Table 3) and the 3 remaining independent alleles are in bold. **A.** Alleles favoring EC. **B.** Alleles preventing EC.

| HLA Alleles |  |  | Meta-analysis |  |
| --- | --- | --- | --- | --- |
| ID | Freq CTR | Freq ECs | OR | P |
| **HLA-B\*57** | **0.03** | **0.15** | **5.92** | **9.482e-43** |
| HLA-C*06 | 0.1 | 0.2 | 2.46 | 1.016e-20 |
| **HLA-B\*27** | **0.04** | **0.09** | **3.21** | **1.89e-17** |
| HLA-A*31 | 0.02 | 0.05 | 2.17 | 1.013e-05 |
| HLA-B*52 | 0.01 | 0.03 | 3.09 | 3.259e-06 |
| HLA-C*01 | 0.04 | 0.07 | 2.04 | 3.847e-06 |
| HLA-C*02 | 0.04 | 0.06 | 1.71 | 0.0002887 |
| HLA-C*14 | 0.01 | 0.02 | 2.1 | 0.002492 |
| HLA-DQA1*02 | 0.14 | 0.2 | 1.46 | 5.669e-05 |
| HLA-DRB1*07 | 0.14 | 0.2 | 1.45 | 6.526e-05 |
| HLA-DQB1*03 | 0.35 | 0.39 | 1.24 | 0.003353 |
| HLA-DPB1*13 | 0.02 | 0.03 | 1.87 | 0.002344 |
| HLA-C*12 | 0.07 | 0.09 | 1.53 | 0.0007661 |

**B.**
| HLA Alleles |  |  | Meta-analysis |  |
| --- | --- | --- | --- | --- |
| ID | Freq CTR | Freq ECs | OR | P |
| HLA-C*07 | 0.31 | 0.17 | 0.46 | 2.942e-17 |
| HLA-B*08 | 0.11 | 0.05 | 0.37 | 7.051e-10 |
| HLA-DRB1*03 | 0.12 | 0.06 | 0.47 | 8.956e-08 |
| HLA-B*07 | 0.12 | 0.06 | 0.51 | 1.11e-06 |
| HLA-DQB1*02 | 0.22 | 0.17 | 0.66 | 1.325e-05 |
| HLA-C*04 | 0.12 | 0.07 | 0.53 | 2.135e-06 |
| **HLA-A\*29** | **0.04** | **0.02** | **0.44** | **0.001139** |
| HLA-B*35 | 0.1 | 0.06 | 0.55 | 2.082e-05 |
| HLA-DQA1*05 | 0.27 | 0.2 | 0.72 | 7.802e-05 |

### LD links between the selected HLA class I alleles and the stepwise regression SNPs

It was interesting to compare the 13 remaining SNPs with the 8 selected HLA class I alleles. Supplementary Table 4A and 4B show the proportion of carriers of the 13 minor SNP alleles among HLA class I carriers (Supplementary Table 4A), and reciprocally, the carriers of the HLA class I alleles among the 13 minor SNP allele carriers (Supplementary Table 4B) in the CTR population. For instance, individuals carrying HLA-B*57:01 almost invariably possess the allele rs111301312 and vice versa (Supplementary Table 4A and 4B). Similarly, 97.5% of the carriers of rs150908530-allele also carry the HLA-A*31 allele while 60% of the HLA-A31 carriers possess the rs150908530-allele, and 40% of the HLA-A*31 carriers harbor the rs9264942 allele (Supplementary Table 4A and 4B). When rs150908530 and rs4418214 were included as covariates, the signal for HLA-A*31 was no longer significant, whereas the signal for rs150908530 remained unchanged when HLA-A*31 was used as a covariate (Supplementary Table 3). Carriers of rs4418214-allele are either HLB-B*27 (48.7%) or HLA-B*57 (48.4%), with carriers of HLA-B*57 and HLA-B*27 always carrying the rs4418214 allele (Supplementary Table 4A and 4B). This suggests that the pronounced effect of rs4418214, which exhibits the highest p-value in Table 1, likely arises from the independent contributions of the HLA-B*27 and HLA-B*57 alleles. Indeed, when HLA-B*27 and HLA-B*57 were included as covariates, the effect of rs4418214 was no longer significant (Supplementary Table 3). Supplementary Table 4A and 4B underscores the profound interconnection between several HLA class I and MHC SNP alleles. By performing regression analyses using either the SNP alleles or the HLA class I alleles as covariates, as previously described, we were able to identify the alleles exerting the most substantial statistical impact (Supplementary Table 3). Following these regression analyses, we ended up finding that HLA-B*52, HLA-A*31, HLA-C*04, HLA-C*07, and HLA-C*14 could be accounted for by SNPs and 3 remaining HLA class I alleles -HLA-B*57, HLA-B*27, HLA-A*29-were likely at the origin of the observed statistical associations. 6 SNPs were also accounted for by covariate analysis with the HLA alleles, and 7 remained after analysis, namely rs9264942, rs150908530, rs1233396, rs79972666, rs9468885, rs1894406 and rs11130312.

Figure 2C presents the final SNP and class I alleles retained after the covariate analysis. Ultimately, we identified 7 SNPs and 3 HLA class I alleles that appear to exert independent statistical effects for EC. In the specific case of rs111301312/HLA-B*57 (R²= 0.98), which was the focus of our previous publication detailing the extensive HLA-B*57 haploblock (12), we presented both variants together in Figure 2C, and counted them solely under the HLA class I allele category in the subsequent analyses.

### Analysis of the most impacting SNPs in CTR and ECs

We analyzed the distribution of the 9 selected SNPs and HLA class I alleles in CTR and EC. As illustrated in Figure 3A, the protective SNPs are more prevalent among ECs, while the two negative SNPs are more frequently observed in CTR individuals. Upon stratifying the ECs into three tertiles based on viral load, we found that the first tertile (CV1), characterized by the lowest viral load, harbors a greater number of protective SNPs compared to the second tertile, which is similar with the third tertile. In the HLA-B*57+ cohort, the EC groups CV1 and CV2 exhibited fewer protective alleles than the CV3 group (Figure 3B), yet retaining a greater number of protective alleles compared to the HLA-B*57+ CTR group. In the cohort of HLA-B*57-subjects (Figure 3C), a different pattern emerged, with the CV1 group of ECs (lowest viral load) carrying more protective alleles (three) than the CV2 and CV3 subgroups (two), whereas HLA-B*57-controls frequently possessed one negative allele, unlike the ECs. Notably, the proportion of protective alleles varied across viral load tertiles for both HLA-B*57- and HLA-B*57+ EC subjects: HLA-B*57-ECs with lower viral loads possessed more protective alleles (Figure 3C), whereas HLA-B*57+ ECs with lower viral load had fewer protective alleles (Figure 3B). This observation suggests that the influence of the B57 allele may obscure the effects of other SNPs for low viral loads, suggesting a distinctive mechanism at play at an early stage of infection for low viral load, which will be further explored in the discussion section.

**Figure 3.**
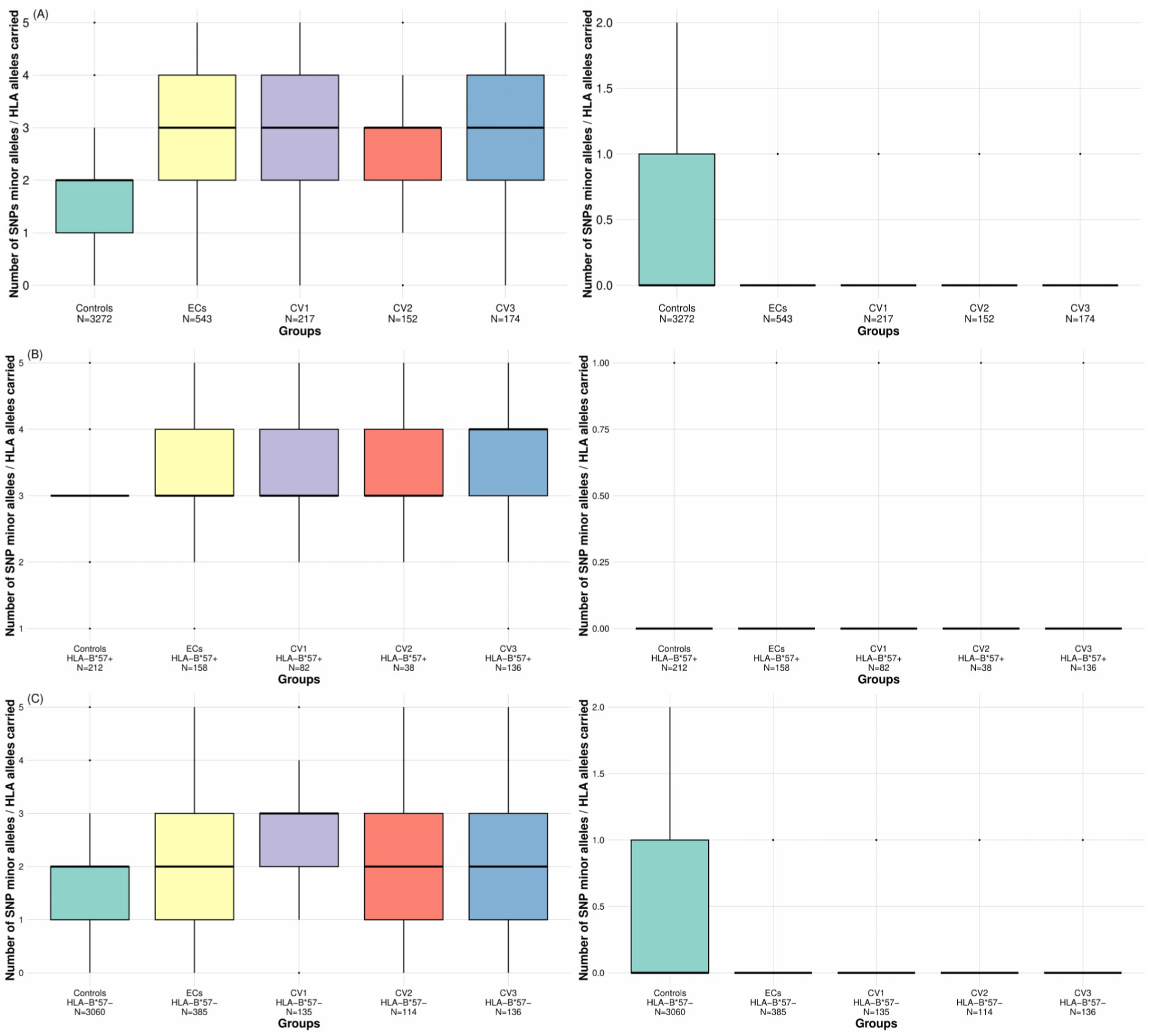
Distribution of protective and negative alleles in Controls, Elite Controllers (ECs), and EC Subgroups Based on Viral Load Tertiles (CV1, CV2, CV3) Boxplot representing the number of minor alleles of the selected SNPs and HLA alleles (7 protective and 2 negative) carried by controls, ECs, and ECs subgroups defined according to the viral load tertiles (CV1, CV2, and CV3 see text). The bold line corresponds to the median. Left panel: presence of the protective alleles. Right panel: presence of negative alleles **A.** Boxplots for the whole CTR and EC groups **B.** Boxplots for the subjects carrying the HLA-B*57 **C.** Boxplots for the subjects not carrying the HLA-B*57 allele.

We then analyzed each SNP in both EC and CTR subjects, as well as within the EC groups stratified by viral load tertiles. The comparison between ECs and CTRs revealed an increase in the minor allele frequency (MAF) of protective alleles in ECs, alongside a decrease in the MAF of negative alleles, consistent with our expectations (Table 1). When examining the MAF among ECs stratified into low (CV1), middle (CV2), and high (CV3) tertiles of viral load, we observed that the MAF remained relatively stable across most SNPs and HLA alleles, with the exception of rs9264942 and HLA-B*57:01 (Table 3). The changes observed among the three groups for rs9264942 mirrored those for HLA-B*57:01, and given that HLA-B*57:01 is in complete linkage disequilibrium (D’=1) with rs9264942, it follows that the enrichment noted for rs9264942 is the full reflection of the enrichment of HLA-B*57:01 (Table 3A). Furthermore, Table 3A indicates a slight decrease in MAF for rs79972666 within the CV1 group, a SNP associated with an overexpression of MICB (log ratio of 0.64, see Table 6). Upon further examination of the subgroups HLA-B*57+ and HLA-B*57-, this decrease predominantly occurs among HLA-B*57+ individuals with low viral load (CV1 group, Table 3B), the subjects who are indeed able to overcome the HIV-induced shedding of MICA and MICB to block NK cells. In HLA-B*57-individuals, there is no modified MAF of rs79972666 in the CV1, CV2, and CV3 groups, probably because the resistance to HIV-1 in the CV1 group of HLA-B*57-subjects is not only solely dependent on NK cell response (Table 3C).

**Table 3.**
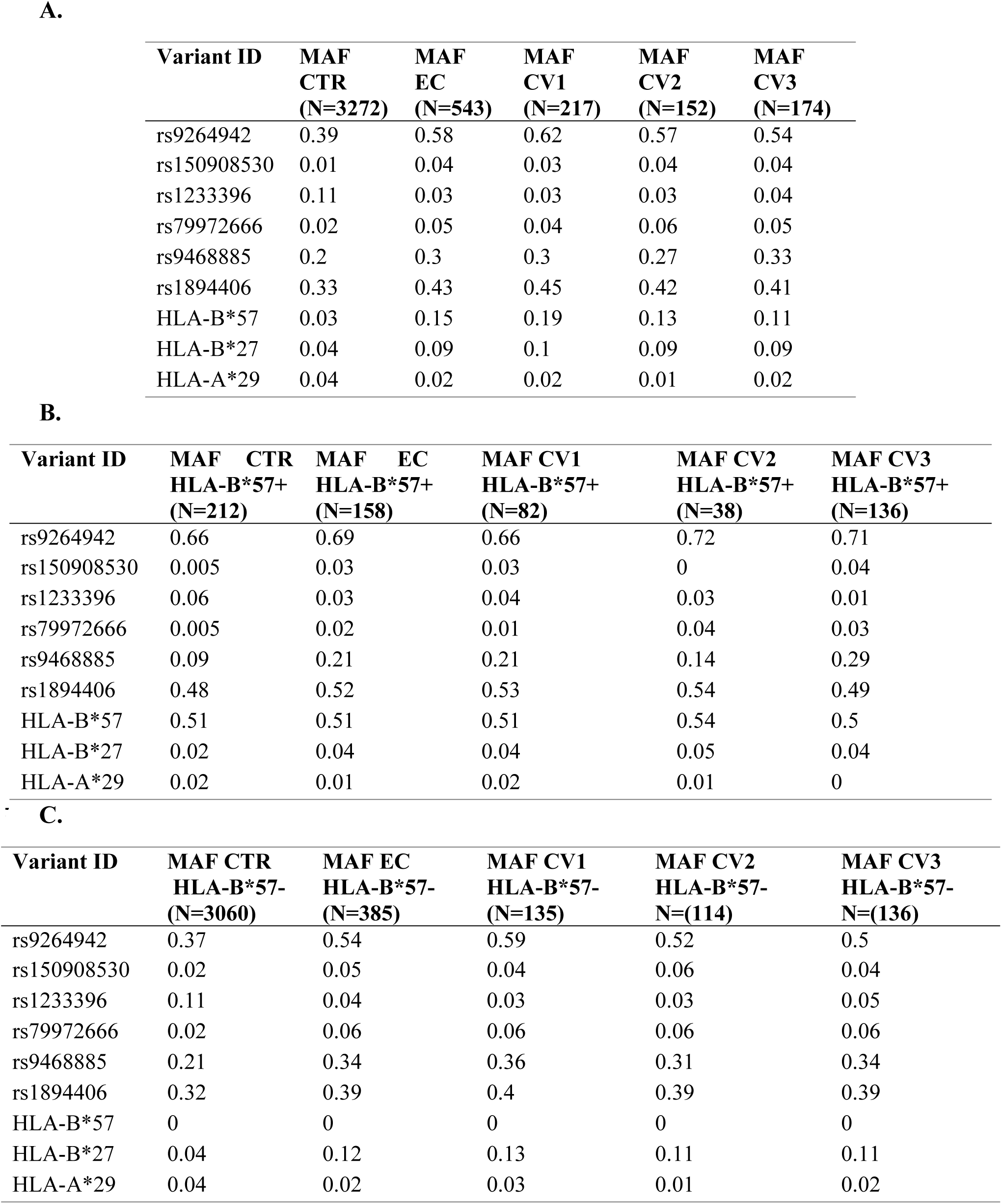
Table showing the MAFs of SNP/HLA alleles in different EC and CTR subgroups. CV1, CV2, and CV3 are the tertiles subgroups of ECs with lower (CV1), middle (CV2), and higher (CV3) viral load. Distribution of the SNP/HLA alleles in A) CTR, EC, CV1, CV2, and CV3 groups B) the HLA-B*57+ subjects of CTR, EC, CV1, CV2, and CV3 C) the HLA-B*57-subjects of CTr EC, CV1, CV2, and CV3.

### Comparison of the LD found in the CTR and in the EC groups

Interestingly, we also built the tree relationships of the selected 6 SNPs and 3 HLA gene variants within the EC subjects (Figure 2D and Supplementary Table 4). By comparing Figure 2C and Figure 2D, one sees that the tree has a quite different shape in the EC population compared to the CTR population, with rs9264942 being positioned at the root of all branches. For instance, 90% of EC carrying rs9468885 minor allele also carry rs9264942, a similar pattern is observed for SNP rs150908530 minor allele. rs1233396 also becomes linked to rs9264942 at more than 80% in the ECs. This suggests that, to harbor a variant associated with a negative impact and be classified as EC, it is also necessary to possess the protective allele rs9264942. HLA-A29, which is decreased in EC (MAF 2% vs 4% in CTR), seems to exert a relatively independent effect from rs9264942 (with 55% carriers in CTR versus 46% in EC). Overall, these observations indicate an increase in specific combinations of SNP/HLA class I alleles (i.e. haplotypes) in the EC carrying the minor allele of rs9264942.

### Haploblock representation of the remaining independent HLA class I and stepwise regression SNPs

In the preceding chapters, we have used stepwise regression and covariate analysis to select the key SNP and classical HLA alleles explaining the main MHC genetic associations with EC. We are aware that the associations found for these SNPs and HLA class I alleles may step from causal SNPs in high LD that have similar association values.

Furthermore, we demonstrated that potential haploblock effects could occur as evidenced for the combined effect of SNP rs9264942 with that of rs111301312/HLA-B*57:01. Such haploblock effects may also be present for the other selected HLA class I alleles akin to those previously described for HLA-B*57:01 (12). Therefore, we decided to use the same strategy and calculated the haploblocks corresponding to the 3 selected HLA class I alleles, encompassing all SNPs whose minor allele contains the HLA class I allele (see methods). Our hypothesis is that a SNP allele within the haploblock may exert a biological effect either concurrently with other SNPs within the haploblock or simultaneously with the HLA class I allele itself, thereby contributing to the observed effect associated with the HLA class I allele.

One might temper this haploblock analysis by asserting that any influential SNP should have been detected through our initial GWAS and stepwise regression processes. Nonetheless, conducting this haploblock analysis serves as an additional safeguard to ensure that no signals have eluded our initial selection strategy. For this reason, we also extended our haploblock analysis by extending it to the 6 remaining SNPs in addition to the three HLA class I alleles. Table 4 presents the haploblocks associated with the selected 6 SNP minor alleles and 3 HLA class I alleles, essentially listing all SNPs whose minor allele “contains” the minor allele of the reference allele. The haploblocks are further summarized in a map that delineates their localization within the MHC region (Figure 4). By construction, the haploblocks of alleles with a lower MAF will contain more SNPs than the haploblocks of SNPs with higher MAFs. Indeed, one can see in Table 4 that there are extensive haploblocks for 2 SNPs and the 3 HLA class I alleles that have a low MAF, namely the haploblock of rs150908530 encompassing 517 kB with 472 SNPs; the haploblock of rs1233396 encompassing 2.8 MB with 1890 SNPs; the haploblock of HLA-B*57:01/rs111301312 encompassing 1.4 MB with 898 SNPs; the haploblock of HLA-B*27 encompassing 319 kb with 355 SNPs; the haploblock of HLA-A*29 encompassing 1.1 MB with 402 SNPs. 4 SNP - rs9264942, rs79972666, rs9468885, rs1894406-exhibit smaller haploblocks and they all have a large MAF except for rs79972666.

**Figure 4.**
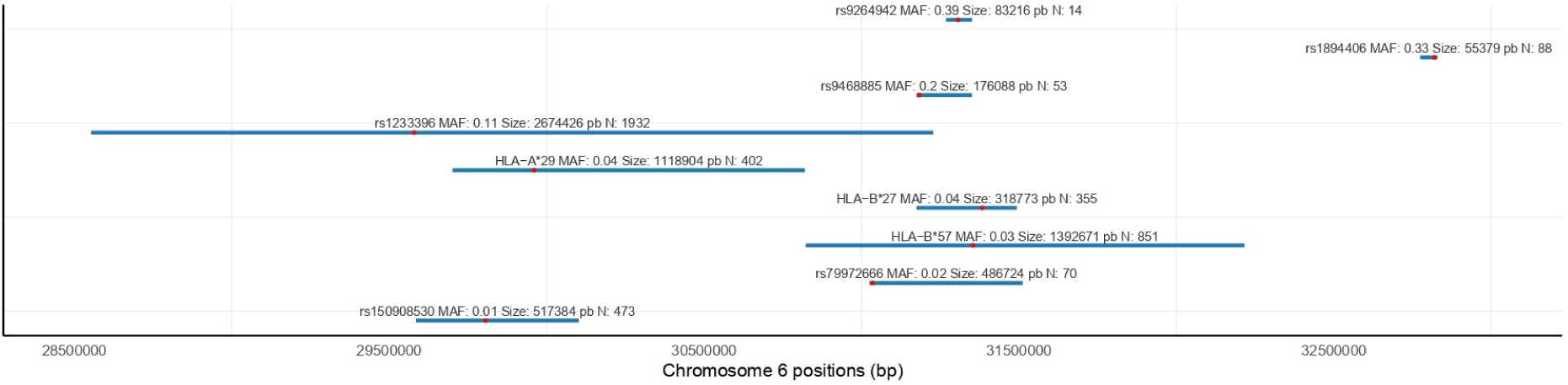
Representation of the haploblocks for the 9 selected SNP/HLA alleles. The blue line shows their extension in the MHC region and the red dot corresponds to the localization of the SNP used to generate the haploblock. The haploblocks are presented from lower MAF (bottom) to higher MAF (top).

**Table 4.** Description of the haploblocks associated with the 9 selected SNP/HLA alleles. The position number corresponds to the (HCG38) version of the human genome. rs111301312 and HLA-B*57:01 are in nearly perfect LD (R²= 0.98), and thus, both variants are treated as a single signal.

| SNP |  | Associated haploblock |  |  |  |
| --- | --- | --- | --- | --- | --- |
| ID | Position | Start | end | size (bp) | N snps |
| rs9264942 | 31306603 | 31268398 | 31351614 | 83216 | 14 |
| rs150908530 | 29806870 | 29585114 | 30102498 | 517384 | 473 |
| rs1233396 | 29579022 | 28553793 | 31228219 | 2674426 | 1932 |
| rs79972666 | 31035668 | 31025467 | 31512191 | 486724 | 70 |
| rs9468885 | 31183636 | 31174520 | 31350608 | 176088 | 53 |
| rs1894406 | 32819279 | 32774141 | 32829520 | 55379 | 88 |
| HLA-B*57/<br>rs111301312 | -- | 30823417 | 32216088 | 1392671 | 851 |
| HLA-B*27 | -- | 31174942 | 31493715 | 318773 | 355 |
| HLA-A*29 | -- | 29701510 | 30820414 | 1118904 | 402 |

The haploblock associated with HLA-B*57:01 exhibits some slight variations compared to our recent publication (12) for two primary reasons: a) our previous study considered only SNPs significantly associated with EC, which accounts for the increased number of SNPs in the current haploblock; and b) we applied more stringent LD criteria here (75% carriers as opposed to 70% carriers), resulting in a smaller haploblock span compared to that reported previously. When two SNPs exhibit high LD (D’ close to 1), the haploblock of one SNP may be included in the haploblock of the other. For instance, rs111301312/HLA-B*57:01 has an LD coefficient D’ equal to 1 with rs9264942. Since rs9264942 is part of the haploblock of rs111301312/HLA-B*57:01, the haploblock derived from rs9264942 haploblock will be included within the haploblock of rs111301312/HLA-B*57:01. Thus, a hierarchical relationship exists among certain haploblocks. These representations are invaluable as they facilitate the investigation of SNP impacts by identifying SNPs that may exert biological effects, present within a haploblock but not in higher order haploblocks within the hierarchy.

### Functional impact associated with the selected SNP and HLA class I alleles

Once the haploblocks built as described above, it was possible to look for the SNP variants exerting a potential biological activity either by protein mutation or by differential gene expression, as we did in our previous work for the HLA-B*57:01 haploblock (12). In our analysis of protein variations, we will take into account all the haploblock variants, but will favor SNPs in high LD with the reference SNP used since the latter have been selected for their highest impact by covariate analysis. We have used ANNOVAR (18) to identify potentially impacting protein variants within the haploblock, and GTEX (19) to identify genes whose expression is potentially impacted by the selected SNP/HLA class I alleles. For sake of simplicity, Table 5 solely presents the protein variations of interest identified in the haploblocks, i.e. satisfying the 2 following criterias : 1. they are deemed damaging according to one of the 2 prediction software Polyphen 2 or SIFT (20,21) the corresponding SNP has a significant p value in the original GWAS. We also examined the SNPs of the haploblocks associated with the 3 HLA alleles when available. In addition, we have provided the complete list of all the protein variations found in the haploblock SNPs in supplementary Table 5. When a protein variant was found in Table 5, we have looked at Genecards (22) to have its function, cell localization, and cell expression, and checked its potential relationship with HIV-1 infection in the scientific literature (see Methods). Similarly, Table 5 presents the impact of the 6 SNP and 3 HLA alleles on differential gene expression, by presenting for each allele, the 3 genes whose expression is mostly impacted in PBMCs according to GTex (see Methods). It is reasonable to consider GTex as a relevant database since ECs have very low viral loads and the transcriptomic profile of their PBMCs is close to that of uninfected subjects (23).

**Table 5.**
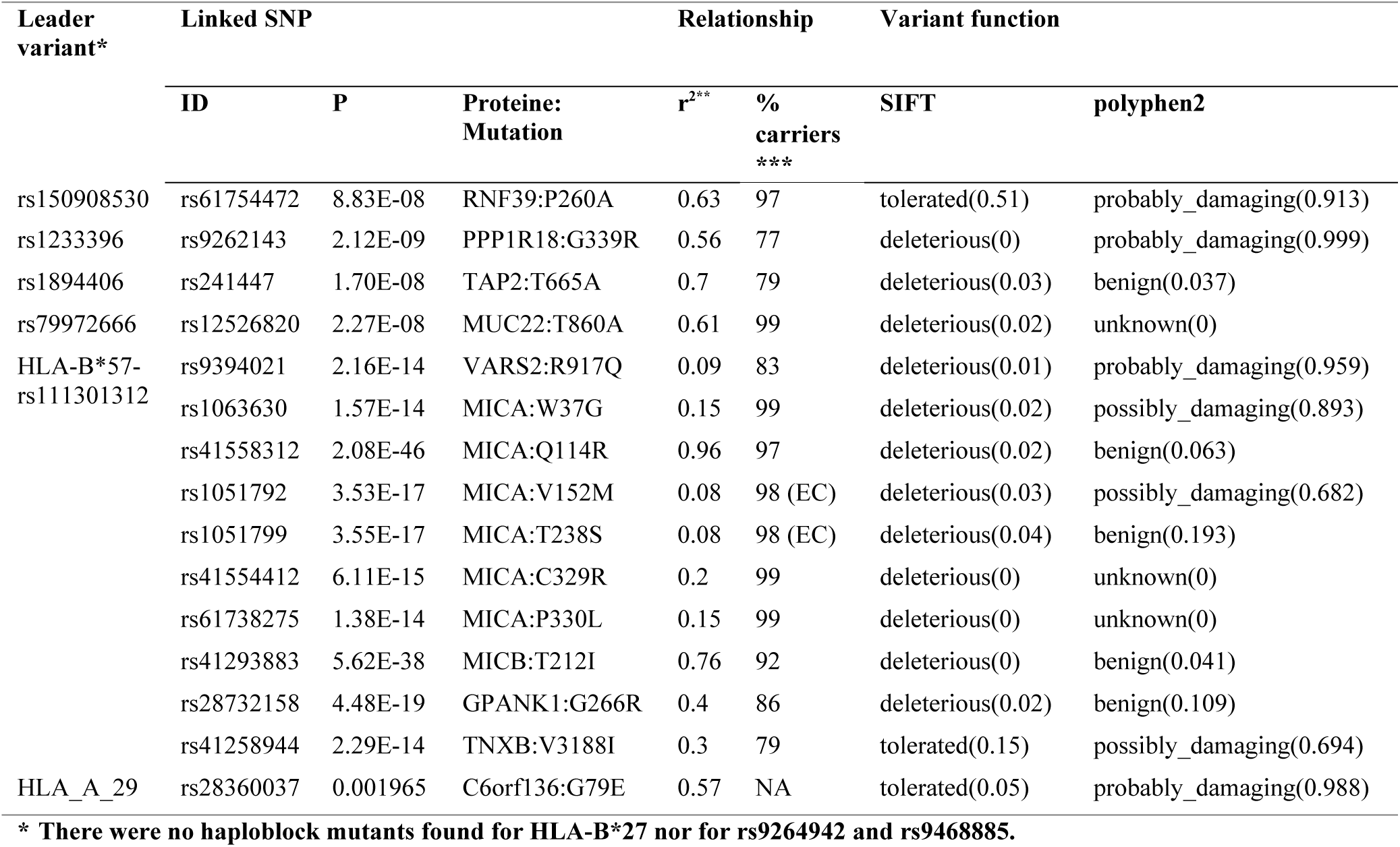
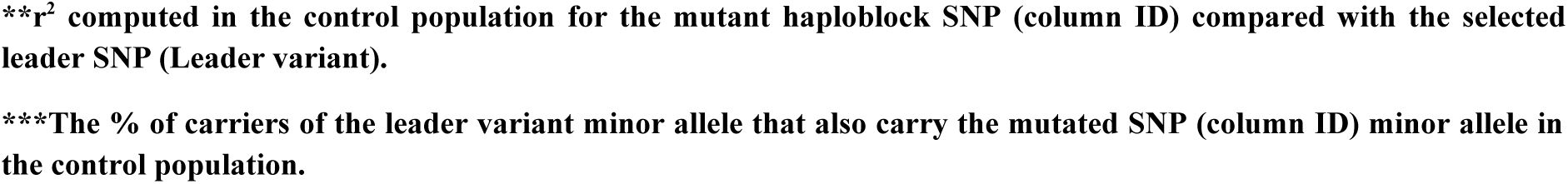
List of the significant SNPs that potentially induce a damaging protein variation in the haploblocks of the 9 selected SNP/HLA alleles.

For SNP rs15908530, there is one protein variant found damaging by Polyphen 2 in RNF39, with a Proline transformed into Alanine at position 260 (P260A, see Table 5). RNF39 is a ring finger protein of size 420 AA, mainly in the nucleus and expressed in all cell types. Two studies have shown that RNF39 knock-down inhibited HIV-1 expression (24,25). This mutation is thus compatible with the protective effect of rs15908530. Interestingly, rs61754472, the SNP associated with this RNF39 mutation has MAF similar to that of rs15908530 in CTR and in EC (with r^2^=0.63). There are also genes with differential expression associated with this SNP: ZNRD1 and HLA-V overexpressed, HCG4P3 underexpressed (Table 6), the two latter being pseudogenes. ZNRD1, a RNA-polymerase 1 subunit of 126 AA, is a nuclear protein present in all cell types. Its down-regulation has been associated with impairment of HIV-1 replication (26,27), but on the opposite, it is not certain that its overexpression will increase HIV-1 replication. Overall, the RNF39 mutation P260A is likely a good candidate to be causal for the EC phenotype.

**Table 6.** Transcriptional impact of the 9 SNP/ HLA alleles in PBMC. List of the 9 selected SNPs/HLA alleles with their transcriptional impact in PBMCs according to GTEx (19) (see Methods). The 3 proteins presented for each SNP correspond to the ones with the 3 best GTEx p values and the effect size correspond to the log of the ratio of gene expression comparing the carriers of the allele (column RS ID) with the non carriers of the allele. The ratio is computed for the SNP allele of column RS ID whose LD (r^2^) with the query variants is indicated in the 3^rd^ column.

| Query | RS ID | $r^2$ | Gene Symbol | Effect Size | P-value |
| --- | --- | --- | --- | --- | --- |
| rs1233396 | rs59113084 | 1 | ZFP57 | 1.07 | 2.33E-31 |
| rs1233396 | rs59113084 | 1 | HCP5B | 0.93 | 2.38E-23 |
| rs1233396 | rs115870917 | 0.99 | HLA-T | -0.64 | 1.25E-10 |
| rs111301312 | rs140220358 | 1 | ZBTB12 | 1.77 | 1.23E-64 |
| rs111301312 | rs41558312 | 0.93 | MICB | -0.94 | 3.74E-34 |
| rs111301312 | rs111301312 | 1 | HLA-B | 0.57 | 6.15E-23 |
| rs150908530 | rs114898901 | 0.86 | ZNRD1 | 0.54 | 9.38E-11 |
| rs150908530 | rs116955874 | 0.93 | HLA-V | 0.9 | 1.53E-07 |
| rs150908530 | rs116997353 | 0.86 | HCG4P3 | -0.62 | 4.78E-06 |
| rs79972666 | rs143179246 | 1 | MICB | 0.64 | 1.07E-07 |
| rs79972666 | rs74558248 | 1 | MIR6891 | 0.63 | 0.0001 |
| rs9468885 | rs1778578420 | 0.97 | XXbac-BPG299F13.17 | -0.16 | 2.21E-09 |
| rs9468885 | rs1778578420 | 0.97 | CCHCR1 | -0.19 | 4.39E-08 |
| rs9468885 | rs1778578420 | 0.97 | HLA-S | 0.33 | 1.27E-06 |
| rs9264942 | rs79709508 | 1 | HLA-S | 0.78 | 3.16E-55 |
| rs9264942 | rs79709508 | 1 | XXbac-BPG248L24.12 | 0.42 | 4.07E-16 |
| rs9264942 | rs79709508 | 1 | ZBTB12 | 0.29 | 6.60E-11 |
| rs1894406 | rs56951180 | 1 | TAP2 | 0.25 | 7.81E-24 |
| rs1894406 | rs56951180 | 1 | HLA-DOB | -0.29 | 4.14E-27 |
| rs3997984 (HLA-B*27) | rs146683910 | 1 | XXbac-BPG181B23.7 | -0.76 | 7.19E-16 |
| rs3997984 (HLA-B*27) | rs114291795 | 0.96 | MICA | -0.32 | 6.28E-05 |
| rs3997984 (HLA-B*27) | rs116666910 | 1 | POU5F1 | 0.28 | 0.0002 |
| rs366030 (HLA-A*29) | rs259948 | 0.89 | HCG18 | 0.53 | 1.44E-14 |
| rs366030 (HLA-A*29) | rs142094780 | 0.89 | XXbac-BPG283O16.9 | -0.38 | 8.83E-09 |
| rs366030 (HLA-A*29) | rs259948 | 0.89 | VARS2 | -0.34 | 1.03E-07 |

SNP rs1233396 is the sole variant preventing EC. In Table 5, we see it is associated with a damaging mutation of PPP1R18 according to both Polyphen 2 (21) and SIFT (20), a glycine replacing an arginine at position 339 (G339R). This protein of 613 AA is well expressed in all cell types and targets Protein Phosphatase 1 to F-actin (28). Importantly, a recent work has shown that HIV-1 prefers low density of cortical actin for virus assembly and particle release (29) suggesting that damaging PPP1R18 might lower F-actin function and thus favor HIV-1 particle production. In Table 6, one can also see that rs1233396 is associated with overexpression of ZFP57 and HCP5B genes, and decreased levels of HLA-T. HCP5B and HLA-T are pseudogenes whose regulatory function is difficult to assess. ZFP57 is a 452 AA nuclear protein involved in DNA methylation and imprinting during development, expressed in most cell types, documented as a transcriptional repressor (30). Overall, the PPP1R18 mutation and the overexpression of ZFP57 seem to be reasonable candidates to prevent the EC phenotype.

SNP rs1894406 is associated with a mutation in TAP2 (T665A). TAP2, a 686 AA protein, is a key component of HLA presentation since it helps load the peptides onto the HLA molecules in the endoplasmic reticulum. The detailed functional structure of TAP2 and TAP1 (its partner for transport) has been explored by Gaudet et al. (31) and does not suggest that the mutation T665A has a real functional effect. In Table 6, one can see that rs1894406 is directly associated with an overexpression of TAP2 and underexpression of HLA-DOB. The overexpression of TAP2 is in line with an improved immune response. HLA-DOB suppresses peptide loading of MHC class II molecules by inhibiting HLA-DM, and should thus reduce the immune response. Overall, the impact of rs1894406 on the overexpression of TAP2 and on the underexpression of HLA-DOB is compatible with its protective effect observed on EC.

SNP rs79972666 is associated with a mutation in MUC22 (T860A). MUC22 is a mucin, a large protein of 1773 AA, produced mainly in the testis and oesophagus. The impact of this mutation on MUC22 function is not clear and the function of MUC22 in HIV-1 infection has not been described. In Table 6, we see that this SNP is associated with overexpression of both the MICB gene and MIR6891, but with relatively weak p values. The role of MIR6891 in HIV-1 infection is not documented in the scientific literature. Overall, it is difficult to affirm any link against HIV-1 infection for this SNP.

SNP rs111301312 which tags HLA-B*57:01 corresponds to a very large haploblock (12) (see Table 4). As shown in Table 5, it is associated with several protein variants. First, a mutation in VARS2 (R777Q) identified as damaging by both Polyphen 2 and SIFT (20,21). VARS2, of size 1063 AA, is a mitochondrial aminoacyl-tRNA synthetase, which catalyzes the attachment of valine to tRNA(Val) for mitochondrial translation and is expressed in all cell types. We have not found any publication proposing a clear mechanism of action for VARS2 in HIV-1 infection. We note that this haploblock SNP, rs939402, has a larger MAF than rs11130312 (17% vs 3%, in CTR). A second protein with variations linked to rs111301312/ HLA-B*57:01 is MICA. MICA, a 373 AA protein, is a stress-induced self antigen, and is also a ligand for the NKG2-D type II membrane receptor on NK cells and binding leads to cell lysis by NK cells. The MICA variations associated with rs111301312/HLA-B*57:01 correspond in fact to a haplotype composed of six simultaneous amino-acid mutations (Table 5): W37G, Q114R, V152M, T238S, C329R, P330L. In addition to these mutations of the haploblock damaging MICA function, a previous work by our group pointed out that HLA-B*57:01 corresponds to the MICA*017 allele which is inactivated and could impact HIV-1 infection possibly by the modification of the MICA cleavage site on the cellular surface (5,32).

Interestingly, another experimental work centered on MICA variants and published in 2016 has also found that MICA*017 presents several isoforms with dysfunctional interactions with NKG2D (33). This MICA allele coincides very well with the SNP rs111301312, since SNP rs41558312 that marks the MICA Q114R damaging mutation in the haplotype is in high LD with rs111301312 (r²=0.96, see Table 5). Explaining how the impairment of MICA function, and thus the prevention of the lysis of infected cells by NK cells, might be helpful for EC is not obvious. However, works published by 2 independent groups, Nolting et al. and Matusali et al. (34,35) provide a quite interesting explanation since they have shown that HIV-1 induces an increased shedding of soluble MICA, which in turn inhibits NK cells and diminishes NKG2D expression. The allele MICA*017 with its damaging mutations (Table 5) would thus not be able to exert an inhibitory effect on NK cells because of the damaging mutations impeding the binding of NKG2D or simply because of the absence of shedding as suggested in our past publication (5). NK cells, which constitute the first line of defense of non specific immunity would thus still be able to lyse and eliminate HIV-1 infected cells at very early stages of infection. The next protein found in Table 5 is MICB which has a function identical to that of MICA. MICB, a 383 AA protein, presents a mutation T180I marked as deleterious by SIFT (20) and associated with HLA-B*57:01. As for the MICA variations, the SNP causing this mutation in MICB, rs41293883, is in high LD with rs111301312 (r^2^=0.76), however it is not clear how this point mutation may impair MICB function since these amino-acids look rather common and SIFT (20) does not see this mutation as damaging. The next protein associated with rs111301312 in Table 5 is GPANK1 with a G266R mutation. GPANK1 is a 356 AA nuclear protein expressed in all cell types, but its function and possible interaction with HIV-1 infection is not known. The last protein of Table 5 associated with rs111301312 is TNXB, with a V3188I mutation. TNXB is a 4244 AA extracellular matrix protein expressed in all cell types, but its function and possible interaction with HIV-1 infection is not known. Other variants of the haploblock of rs111301312 are in CCHCR1, a protein involved in mRNA metabolism, and could have a potential impact since they correspond to mutations of cysteines, but are not presented in Table 5 since the SNPs are not significant (Supplementary Table 5).

Regarding differential gene expression (Table 6), rs111301312 is associated with the overexpression of HLA-B and ZBTB12 with rather high log ratios (resp. 0.57 and 1.77). ZBTB12 is a transcriptional factor of 459 residues, expressed in all cell types, with a role in the regulation of human endogenous retroviruses, particularly in the upregulation of HERV-K (36). Interestingly, a publication has shown in vitro that HERV-K may interfere with HIV-1 infection and lead to a progeny of less infectious HIV-1 particles (37) providing a possible explanation for the impact of ZBTB12 overexpression. Rs111301312 is also associated with the decreased gene expression of MICB, thus possibly limiting the retro-inhibitory effect of shedded MICB proteins induced by HIV-1 infection (34,35) and favoring the early NK cell response.

Overall, the haplotypic mutations impairing MICA function seem to be compatible with the control of HIV-1 infection through NK cells at an early stage of infection. The roles of VARS2, GPANK1, or TNXB in HIV-1 infection have not been described in the literature. The overexpression of ZBTB12 and underexpression of MICB seem compatible with the control of HIV-1 infection. Of course, the role of HLA-B*57:01 as protein presenting peptides for the immune system could also be an explanation for the contribution of rs111301312 for EC, once the infection is well set.

HLA-A*29 presents a large haploblock and we could look at his potentially impacting variants in Table 5. We find again a mutation of the 420 AA protein RNF-39, D268N detected by Polyphen 2. There is no clue to understand how this point mutation may impact RNF39 function since D and N are close amino-acids, and this mutation is clearly less convincing than the P260A identified for rs15908530. The second variant found for HLA-A*29 deals with a variant (G79E) in c6orf136 which is an integral membrane protein of 315 AA protein expressed in all cell types. There is no clear link of this protein with HIV-1 infection in the bibliography.

For HLA-A*29 the gene expression profiles does not present very low p values, except for HCG18. It is a lncRNA which has been related to colorectal cancer but not to AIDS. It may be effective in AIDS but it is difficult to provide an interpretation through the scientific literature, in spite of the relatively high expression log ratio of 0.53 (Table 6).

There was no significant variant described for rs9264942 and HLA-B*27 in Table 5. Of course HLA-B*27 allele itself is a good candidate as protein presenting peptides to the immune response.

In Table 6, rs9264942 is associated with the overexpression of HLA-S with a relatively high log ratio (0.78), and with XXbac-BPG248L24,12 and ZBTB12. HLA-S is a pseudogene with no described specific relationship with HIV-1 infection, and there is neither any relationship with HIV-1 infection for XXbac-BPG248L24,12. We retrieve also ZBTB12 also found for rs111301312, with a much lower expression coefficient (0.29).

For HLA-B*27, in terms of gene expression (Table 6), we found underexpression of XXbac-BPG181B23.7 with unknown function, underexpression of MICA but with a rather weak log ratio (−0.32) compared to the MICB underexpression of HLA-B*57:01 (log ratio = −0.94), and overexpression of POU5F1 with no significance on AIDS literature.

## Discussion

### General observations on the work performed

In this work, we analyzed the GWAS data from the ICGH project to identify genetic associations with EC. We identified a substantial number of genome-wide significant associations, totaling 2,626, all localized within the MHC region. When comparing this result to the previous study of Pereyra et al (7), we find many more genome-wide significant signals (p<5.10^-8^), i.e. 2626 versus 323. This discrepancy can likely be attributed to the more extreme phenotype studied (EC vs viral controllers) and possibly by the progress in the bioinformatics database. The high number of significant associations (2,626) relative to typical GWAS results and the number of SNPs in the MHC region (31,500) is probably due to the extensive LD existing between SNPs in the MHC region.

In order to break this complexity and gain a clearer understanding of the potential causal SNPs and underlying biological mechanisms, we employed a combination of stepwise regression on all SNPs, integrated the association results for HLA class I and class II gene alleles, and performed LD analysis with covariate adjustments to eliminate signals statistically dependent on others due to LD. Our final analysis yielded six SNPs and three HLA class I alleles (Figure 2C): rs9264942, rs150908530, rs1233396, rs79972666, rs9468885, rs1894406, HLA-B*57:01, HLA-B*27, and HLA-A*29, for which we constructed their haploblocks. They represent a synthesis of the statistical effects observed in the initial 2,626 signals, demonstrating our success in reducing the complexity of the original GWAS.

Several key observations emerged from our study. Firstly, the haploblock associated with rs111301312 (a tag for HLA-B*57:01) was of particular interest since in the original GWAS, the OR for this SNP was 6, but it decreased to 2.35 after stepwise regression (Table 1). When testing SNPs within the haploblock as covariates in the association analysis, we found that SNP rs9264942 was the primary cause of this decrease. This finding demonstrates that the strong association observed for HLA-B*57:01 is partly due to the effect of rs9264942, corroborating our previous work that identified additional functional SNPs within the large haploblock of HLA-B*57:01.

Secondly, we observed that several HLA class I and II alleles lost their significance when other HLA alleles were used as covariates in the association analysis (Figure 2B and Supplementary Table 3). This confirms that many alleles across various HLA class I and II genes are linked through LD, a phenomenon well-documented in ancestral haplotypes involving HLA-A, HLA-B, HLA-C, and HLA-DR/DP/DQ genes (15,16).

Thirdly, we found that many signals associated with HLA class I and II alleles were actually driven by combinations of certain MHC SNPs (Supplementary Table 3). In other words, we demonstrated that several HLA class I and II associations can be attributed to the LD between these HLA alleles and MHC SNPs involving other genes. This observation is crucial for all studies investigating associations with HLA class I and II alleles. When identifying associations with HLA class I or 2 alleles, it is essential to consider potential causal associations with other MHC SNPs in LD, as this can reveal new mechanisms of action, as demonstrated in the present study.

These results are important since they show for the first time that the MHC effect for EC is not solely attributable to the HLA class I and class II alleles, but also to SNPs present in other MHC genes. In continuation of our previous work on HLA-B*57, our current study simultaneously examined the roles of class I and class II alleles and other MHC gene variants. Our results indicate that several significant SNPs are actually driven by HLA class I alleles, and several significant HLA class I and II gene alleles are driven by SNPs (Figure 2B and Supplementary Table 3). We have clearly demonstrated that HLA class I and II gene alleles are not the sole drivers of the EC phenotype.

### Main results and biological interpretation

With our protocol, we have identified 9 main variants (SNP and HLA class I alleles) summarizing the 2,626 statistical associations found with EC. We acknowledge that some information may have been lost during our analysis, but the primary independent signals should still be there thanks to the exhaustive protocol followed. During the stepwise regression step, SNPs in high LD could not be distinguished, and one SNP among them was randomly selected. This implies that the underlying biological cause for each of the selected SNP/HLA alleles should rather be SNPs in high LD with them (let’s say r^2^>0.5). Once the SNP and HLA alleles were selected, to avoid missing any biological information, we examined not only protein variants among SNPs in high LD but also all SNPs within the haploblocks associated with these 9 variants, to ensure no signals were missed. This broadened screening also served to confirm the quality of our protocol by verifying if the biologically impactful SNPs identified were indeed SNPs in high LD with the 9 identified variants.

Of course, the three identified HLA class I alleles have a known biological role in peptide presentation to the immune system, but it was also intriguing to investigate other MHC SNPs in high LD with potential biological impacts. Using Annovar (18) (Table 5) and GTEx (19) (Table 6), we extensively looked at all the protein variants, especially the potentially deleterious ones impacting protein function, and at the mRNA differential expression in PBMCs that could significantly impact the EC phenotype. From the identification of these variants, it became thus possible to make hypotheses on how they could impact the EC phenotype.

In Table 5 (analysis of the protein variants), several variants of interest were found among the 9 SNP and HLA allele haploblocks, leading to potentially impaired proteins according to SIFT or PolyPhen-2 (20,21). After detailed analysis and evaluation of their potential link with HIV-1 infection (see Section “Functional impact…”), three mutations appear particularly relevant: the P260A mutation in RNF39, whose knockdown has been associated with blocking HIV-1 replication in two experimental studies (24,25), the G339R mutation of PPP1R18 that acts on F-actin which inhibits viral budding (28,29) and the MICA*017 mutant allele (5) with an impaired function (mutation and/or absence of shedding) that should limit the blockade of NK cells caused by the MICA/B shedding induced by HIV-1 infection (34,35). It is noteworthy that the variants corresponding to these mutations were in high LD with the initially selected SNPs: the RNF39 mutation corresponds to SNP rs61754472 in high LD with rs150908530 (r^2^=0.63), the PPP1R18 variant corresponds to SNP rs9262143 in high LD with rs1233396 (r^2^=0.56), and the MICA*017 allele corresponds to SNP rs41558312 in high LD with rs111301312 (r^2^=0.96). We do not exclude a possible direct effect of the HLA class I B*57:01 in association with EC. HLA-A*29 is associated with a G79E mutation in C6orf136, with no assigned biological explanation, but its negative effect on EC could be mediated by its class I peptide presentation function. Similarly, there was no significant mutation in the HLA-B*27 haploblock, and its positive effect on EC could also be mediated through its peptide presentation function to the immune system.

In Table 5, there is inherently a high LD between a SNP inducing the differential expression of genes and our identified SNP/class I alleles since GTex only works with SNPs in high LD (see Methods). The detailed analysis of Table 5 made in the previous section has yielded highly relevant information for all SNPs. A global analysis shows that SNPs favoring EC are all associated with increased expression of HLA genes and pseudogenes (HLA-B, HLA-V, and HLA-S), while rs1233396, which prevents EC, is associated with decreased expression of HLA-T. This concordance suggests an effective role for these HLA genes even though three of them are pseudogenes. Additionally, the GTEx analysis highlighted several new genes such as ZFP57 (rs1233396), ZBTB12 (rs111301312 with its high log ratio of 1.77), TAP2, HLA-DOB (rs1894406), and MICB (rs79972666 and rs111301312). We have seen that ZFP57 is a 452 AA protein involved in DNA methylation and imprinting during development active in adult PBMCs (30), ZBTB12 is a 459AA transcriptional factor known to regulate the human endogenous retroviruses and possibly limit HIV replication (36,37), MICB is a major factor involved in NK cell lysis, TAP2 and HLA-DOB are associated with the peptide presentation by the immune system. In the case of HLA-B*57:01, the decreased expression of MICB aligns with the known dysfunction of the MICA*017 allele, by limiting MICB shedding and its inhibitory effect on NK cells (34,35). We will discuss later the apparent discrepancy in MICB expression between HLA-B*57:01 (log ratio −0.94) and rs79972666 (log ratio 0.64), while both favor EC. Another interesting observation is the strong genetic effect of rs9264942, which translates into a rather weak overexpression of ZBTB12 (log ratio 0.29) and a more significant overexpression of HLA-S (log ratio 0.78), underscoring the likely importance of HLA gene expression in the EC phenotype. Overall, we have identified several biological mechanisms potentially contributing to the EC phenotype in HIV-1 infection. Among them, we put forward the very convincing multi-action mechanism linked of HLA-B*57:01 involving first the NK cell response already known to be important in AIDS (38), and a surprisingly strong overexpression of ZBTB12 that could impair HIV-1 progeny infectivity via HERV-K (36).

### Comparison with our previous work on the HLA-B*57:01 haplotype

In this study, we have performed a comprehensive analysis of all MHC signals associated with EC, whereas our previous study (12) focused exclusively on the HLA-B*57:01 haploblock. When comparing the results of both studies, we observe a high degree of similarity in the characterization of the HLA-B*57:01 haploblock and in the differentially expressed genes, and slightly less congruence in the description of the protein-impacting mutations. Indeed, in this study, we applied more stringent thresholds to emphasize the major signals. Consequently, we observed that the newly delineated HLA-B*57:01 haploblock is smaller in Kb since we used here the criteria of 75% carriers instead of using 70% employed in our previous study (see Methods). Nevertheless, it encompasses a greater number of SNPs since we did not restrict our analysis to SNPs that were statistically significant in the GWAS. Regarding the differential gene expression analysis, in our previous work (12), we had considered all SNPs within the HLA-B*57:01 haploblock, ultimately zooming on the genes MICB, HLA-B, and ZBTB12 which were common to both European and African-American populations. In this study, we indeed identified these 3 genes as directly associated with HLA-B*57:01 by GTex, and extended the study to the genes differentially expressed associated with the other selected SNPs/HLA class I alleles. The majority of differentially expressed genes are associated with immune functions, including the HLA-X (X=B,S,T,V) genes and TAP2. Regarding protein variations, we have focused here on inactivating mutations, specifically insisting on the damaging MICA mutation (allele MICA*017) not recognized in our recent study (12) but in a more ancient study by our group (5), and that could explain the mechanism of escape from HIV-induced NK cell blockade described by several groups (34,35).We have also noted the interest of the PPP1R18 mutation associated with rs1233396. In our earlier HLA-B*57:01 publication (12), we had highlighted all non-synonymous mutations present in the haploblock, some of them being of interest due to the protein functions (e.g., DXO, NOTCH4), although these were not classified as damaging (unlike the MICA mutation) and were not unique to HLA-B*57:01, as we had examined all non-synonymous mutations within the haploblock.

In our previous publication (12), we had noted the specific enrichment of 2 SNPs in EC with lowest viral load, namely HLA-B*57:01 and rs4418214. Here, we could explain why the second SNP rs4418214 is also enriched in EC with lowest viral load: the minor allele of this SNP exactly encompasses both HLA-B*57:01 and HLA-B*27. As a consequence, when there is an enrichment of HLA-B57 individuals among EC with low viral load, there will also be an enrichment of subjects carrying the minor allele of rs4418214. As shown in Table 3, there is indeed a significant enrichment of HLA-B*57:01 and HLA-B*27 individuals among EC subjects compared to uninfected controls, but this enrichment is even more pronounced at lower viral loads only for HLA-B*57:01 (Table 3A). This enrichment in EC with lower viral load does not extend to any of the other SNP/HLA alleles selected in our study, except for the passenger effect of SNPs in high linkage disequilibrium with HLA-B*57:01 (such as rs9264942, rs1894406).

### Why not all HLA-B*57:01 subjects are ECs ?

In our previous work, we noted that the protective effect of HLA-B*57:01 was effective even in small cohorts with virtually no EC (12). This suggests that the statistical correlation between HLA-B*57:01 and viral load at setpoint arises from the entire population of HLA-B*57+ subjects, gradually diminishing over time among these individuals. If this association did not decline, all HLA-B*57+ individuals would remain non-progressors. The viral load at infection will certainly be an impacting factor as we have observed an increase in the proportion of HLA-B*57+ subjects among ECs with lowest viral load (CV1 group) : this proportion of HLA-B*57+ subjects rises from 6% in CTR, to 20% in regular ECs, and up to 40% in the EC CV1 subgroup characterized by very low viral load (Table 3 presents only the MAF). This shift from 6% to 40% of carriers in the CV1 EC subgroup indicates that the HLA-B*57:01 allele is a major determinant for the status of EC with low viral load. But again, not all HLA-B*57+ subjects are ECs. In our previous study (12), we sought gene variants present in HLA-B*57+ ECs that were absent in HLA-B*57+ CTRs, focusing exclusively on variants enriched in ECs with low viral loads compared to those with higher viral loads. These latter variants are in fact the ones present in the HLA-B*57:01 haploblock which remains structurally unchanged in both ECs and CTRs (12). In the present study, we have examined the presence of six additional SNP/HLA alleles not linked to HLA-B*57+ (all except HLA-B*57:01, rs9264942, and rs666) in HLA-B*57+ individuals, comparing HLA-B*57+ CTR with HLA-B*57+ EC for these six SNPs (Figure 3B). Unsurprisingly, Figure 3B indicates that HLA-B*57+ ECs possess a significantly greater number of these six alleles compared to HLA-B*57+ CTRs, providing insights into why not all HLA-B*57+ individuals are ECs. Furthermore, other factors such as the initial infectious dose, additional rare variants, the interferon response, or HIV restriction factors may also contribute to the EC status. In Figure 3B, we observe that HLA-B*57+ ECs with low viral load carry fewer protective alleles than those with higher viral loads, while the opposite trend is observed in HLA-B*57-EC individuals. We can hypothesize that HLA-B*57+ EC subjects exhibiting very low viral loads at setpoint likely contracted the virus through minimal exposure, and their NK cell defenses, along with other anti-HIV-1 mechanisms (e.g., ZBTB12), are sufficient to control the virus and maintain low viral loads. Conversely, HLA-B*57+ EC individuals with higher viral loads continue to manage the virus but may require additional protective alleles to regulate viremia, as evidenced by comparisons with HLA-B*57+ CTRs who exhibit fewer protective alleles (Figure 3B). In contrast, HLA-B*57-subjects are likely to progress if they lack the NK cell defenses (neutralized by the shedding of MICA/MICB) or the cellular anti-HIV defenses (such as ZBTB12). It is reasonable to suspect that all individuals (HLA-B*57+ and HLA-B*57-) have an equal opportunity to acquire other protective factors (either the genetic ones identified here, or others linked with interferon response, viral restriction factors etc..), this results in 2 consequences : a. HLA-B*57-ECs present in the EC CV1 group (very low viral load) may lack an initial functioning NK cell defense (34,35) but require the presence of additional protective alleles as illustrated in Figure 3C and b. HLA-B*57+ individuals are likely to be enriched among those initially infected with low viral loads, as the majority of HLA-B*57-individuals are expected to progress, except for the rare cases possessing several additional protective factors. When we assess the average number of protective alleles in Figure 3, we find that the median number of protective alleles for HLA-B*57+ individuals in the EC CV3 subgroup reaches 4, indicating a median of 3 additional alleles beyond HLA-B*57:01 (Figure 3B). Similarly, the HLA-B*57-EC subjects in the CV3 subgroup also exhibit a median of 3 protective alleles in Figure 3C. The boxplots for both HLA-B*57- and HLA-B*57+ subjects remain higher than those of their respective controls, confirming the protective impact of the alleles identified in this study. This observation leads to several conclusions: first, as discussed before, additional factors of protection exist beyond the genetic alleles selected in this study, as several ECs exhibit 2 or fewer protective alleles, similar to the controls. But this result also suggests that in the EC CV3 subgroup, HLA-B*57:01 itself does not confer as much protection for EC CV3 subgroup compared to the other selected alleles, otherwise the median of Figure 3B for HLA-B*57+ subjects would be 3 protective alleles as in the HLA-B*57-subgroup (Figure 3C), rather than the 4 protective alleles observed in Figure 3B. This supports the interpretation that the HLA-B*57:01 effect for EC occurs early in the course of infection, reinforcing the notion that all HLA-B*57+ subjects experience an early advantage regarding EC status, likely due to the non-blockade of the early NK cell response and the intra-cellular defense mechanisms mediated by ZBTB12.

### Natural history of infection and molecular etiology of the resistance to HIV-1 disease progression

Figure 3 illustrates that the median number of protective SNP/HLA class I alleles carried by EC individuals is three, a threshold rarely met in control subjects (CTR). However, approximately 25% of EC individuals possess only two protective markers, indicating that additional factors may influence the EC status (low infectious dose, rare genetic variants, presence of anti-IFN antibodies which have been shown to be critical in COVID-19 infections (39), HIV-restriction factors (40)…). These additional factors were not explicitly considered in our genetic analysis but we can consider that they may randomly apply to all ECs, independently of their genetic background.

In the preceding paragraph, the results have led us to hypothesize that the effect of HLA-B*57:01 on EC status manifests early and may be explained through two mechanisms: an effective NK cell defense and the potential intra-cellular protection conferred by ZBTB12 which may influence non-infectious viral progeny via HERV-K. Upon infection, an individual may encounter either a low or high viral dose. In case of a low dose, it is plausible that the virus will gradually spread in the body, unless the individual is HLA-B*57+ and can control the virus through one of the aforementioned mechanisms, or alternatively possesses multiple protective alleles, as illustrated in Figure 3B and Table 3C, where carriers of the minor alleles of rs9264942, rs9468885, and HLA-B*27 appear to be more prevalent among HLA-B*57-subjects with low viral load (CV1 group). If a subject gets infected with a high dose, the virus will colonize multiple sites within the body and it will be harder to control the infection. In such case, the control of viral load by HLA-B*57+ subjects will require additional protective mechanisms compared to the CTR group as seen in Figure 3C with for instance more rs9264942 and rs9468885 minor allele carriers in the HLA-B*57+ CV3 group (Table 3B), and for the HLA-B*57-CV3 group we also observe additional protective alleles compared to HLA-B*57-CTR (Figure 3B and Table 3C).

From very low viral load, there will be a progression towards higher viral load for all HLA alleles except for HLA-B*57:01 explaining the enrichment of the latter in the low viral load EC CV1 subgroup (Figure 3B and Table 3). In cases of larger initial viral loads, the presence of multiple protective alleles may enhance resistance to the progression of infection, potentially in conjunction with additional factors (restriction factors, interferon responses, etc..) that provide stochastic protection against HIV-1 dissemination.

Interestingly, the SNP rs79972666 is associated with overexpression of MICB (Table 5), and based on our hypothesis, this may hinder NK cell activity during early stages of infection. Indeed, Table 3B reveals a lower frequency of rs79972666 among HLA-B*57+ ECs in the CV1 group, suggesting a potential detrimental impact on the NK cell defense mechanism in HLA-B*57+ individuals. A final noteworthy consideration that underscores the significance of MICB in HIV infection is that we observe in Table 5 an overexpression of all HLA genes associated with SNPs that favor elite control. Given that MICB is an ancestral form of HLA molecule with similar function (albeit without peptide presentation), it is reasonable to expect similar patterns of protection. It is indeed well-established that NK cells play a protective role following primary infection (38). In summary, while MICB overexpression may not confer protection during the initial stages of infection with low viral load due to HIV-induced shedding and NK cell blockade (34,35), it may provide protective effects at later stages. This is similarly true for MHC class I alleles, such as HLA-B*57:01 and HLA-B*27, which have been documented for their protective effects against HIV-1 infection, extending beyond elite controllers (10).

## Conclusion

This study offers significant insights into longstanding questions regarding elite control in HIV-1 infection, particularly the enrichment of HLA-B*57:01 among elite controllers and the reasons why not all HLA-B*57+ individuals are elite controllers. We have compiled all the genetic variants associated with EC, found the ones potentially impacting protein function (through mutation or through differential expression) and analyzed their distribution according to the viral load (CV1, CV2, CV3 subgroups). We have then tried to put the pieces of the puzzle back together by taking into account the published information regarding the protein functions, their interaction with HIV-1, and the physiopathogenesis of HIV-1 infection.

Our findings related to HLA-B*57:01 are particularly noteworthy, as they strongly suggest an association between this allele and a reduced natural killer (NK) cell activity characterized by a mutated MICA allele and decreased MICB expression, along with increased production of HERV-K possibly interfering with HIV-1 multiplication. It may appear somewhat paradoxical that the diminished function of MICA/MICB observed in HLA-B*57:01 carriers may be the very reason their NK cell activity can overcome the HIV-1 evasion mechanism that involves the shedding of these molecules in the environment of infected cells (34,35). It is important to note that the NK cell function of HLA-B*57:01 carriers is not entirely abrogated, as there is still some expression of MICB (and additionally, a second chromosome 6), but biological phenomena are often quantitative. From this standpoint, we may hypothesize that, for certain diseases, a population-level statistical impact could be observed due to this less functional NK cell response in HLA-B*57:01 carriers. Indeed, HLA-B57 presents the strongest association with psoriasis, a highly prevalent dermatological condition, and most interestingly, NK cells appear to be associated to this pathology in affected patients (41). Interestingly, the HLA-B*57:01 haploblock also contains SNP alleles with larger MAF corresponding to inactivating mutations in the protein CCHCR1 (Supplementary Table 5) described as a contributor to the physiopathogenesis of psoriasis (42). But the strongest association in psoriasis remains with HLA-B*57:01 (41) confirming the specific impact of this haplotype.

Our research underscores the multifaceted role of the MHC region, central command of the body and cell defense systems, in combating HIV-1 infection, extending beyond the traditional class I and II antigen presentation pathways. For the first time, we highlight defense mechanisms associated with HLA-B*57:01 that contribute to low viral load, which directly involve MICA/MICB and potentially ZBTB12 genes for early immune responses, alongside the contributions of other variants impacting other genes (TAP2, DOB, RNF-39 etc..), including PPP1R18 which may inhibit viral budding throughout the infection process. Conversely, the virus has evidently evolved quite sophisticated mechanisms to evade this MHC central command pressure, albeit such strategies appear less effective in front of the HLA-B*57:01 haploblock. More generally, it is essential to recognize that associations observed with HLA class I and II molecules may represent only a fraction of the underlying complexity present within this sensitive gene-rich MHC region.

## Materials and Methods

### Participant Phenotypes and Case/Control Matching

In this study, we used genotyped data from the International Collaboration on HIV-1 Genomics (ICGH). The ICGH project, initiated in 2012 with the support of the National Institutes of Health, aimed to consolidate genomic datasets from HIV-1 infected individuals worldwide. The consortium comprised 26 cohorts of seropositive subjects genotyped on diverse platforms, representing four continents (US, Europe, Australia, Africa). Genotypes for uninfected control individuals were obtained from three participating centers, the Illumina genotype control database and the Myocardial Infarction Genetics Consortium (MIGen) (43,44). Each dataset underwent preliminary quality control procedures before centralizing all the data for combined analysis. However, to ensure consistency, additional quality control measures were implemented once the data were submitted, as described in the initial publication of ICGH (44).

All the individual cohorts contributing to the ICGH effort obtained ethical approval from their respective country institutions. The initial publication of ICGH in 2013 focused on susceptibility to HIV-1 infection (44) and included a comparison of 6,300 seropositive individuals with 7,200 uninfected controls of European descent. A subsequent publication in 2015 investigated the genetic association with viral load setpoint among the 6,300 seropositive subjects of European descent (45). In these studies, the cohorts were categorized into six groups of matched cases and controls based on genotyping platforms and geographic origin.

The present study focuses on two groups of Elite Controllers (EC) from the European cohorts, characterized by a viral load of less than 1,000 copies/mm3. Sufficient cases were available for our analysis in ill1 (418 ECs) and ill2 (125 ECs). Corresponding uninfected matched controls included 2,759 subjects in CTR1 and 513 subjects in CTR2. The matching process was described in the initial ICGH publication (44).

For our analyses, we categorized the ECs into three subgroups based on their viral load: CV1 with viral loads between 1 and 100 copies/mm³ (217 ECs), CV2 with 100 to 400 copies/mm³ (152 ECs), and CV3 with 400 to 1,000 copies/mm³ (174 ECs).

### Participant genotypes

The participants of European descent within the ICGH consortium were organized into matched case-control groups using a two-stage case/control matching strategy, as outlined in the initial publication of ICGH (44). This resulted in four clusters: Group 1 consisted of participants from the Netherlands genotyped on the Illumina platform, Group 2 included participants from France genotyped on the Illumina platform, Groups 3 and 4 encompassed participants from North America and non-Dutch/non-French European regions genotyped on the Illumina platform, and Groups 5 and 6 comprised participants from North America and non-Dutch/non-French European regions genotyped on the Affymetrix platform.

The ill1 EC group and its matching controls (CTR1) originated from Group 3, while the ill2 EC group and its matching controls (CTR2) were derived from Group 4. Genotypic data for these groups were generated using three genotyping arrays: Illumina 550, Illumina 650, and Illumina 1M. Prior to analysis, we conducted standard quality control (QC) procedures on each set (CTR1/ill1 and CTR2/ill2) to ensure the use of clean data.

Specifically, we filtered out monomorphic or rare variants with a minor allele frequency (MAF) less than 1%, as well as structural variations such as insertion-deletions. We also excluded sites with a missingness rate above 0.02 or a Hardy-Weinberg equilibrium (HWE) p-value lower than 10-6. Moreover, we examined the consistency between recorded sex information and sex inferred from the X chromosome, assessed heterozygosity and homozygosity rates, and verified the level of relatedness by calculating identity by descent (IBD).

### SNP Imputation

Before imputation, additional quality control (QC) steps were applied to the two sets of European descent (ill1/CTR1 and ill2/CTR2). These QC measures utilized the checkbim steps of the McCarthy Group Tools (46), which involved removing SNPs with A/T and G/C alleles if the minor allele frequency (MAF) exceeded 40%. SNPs with discordant alleles, those with more than a 20% difference in allele frequency, and SNPs not present in the 1000 Genomes reference panel were also excluded. These QC steps ensured the utilization of high-quality data for the subsequent imputation process, enabling accurate results.

For ill1/CTR1, a total of 387,495 common SNPs were imputed, while for ill2/CTR2, 271,572 common SNPs were imputed. The imputation process followed a specific protocol: phasing was performed using Eagle2.4 (47), and imputation was carried out on the TOPMed Imputation Server (48) using Minimac4 (49) and the TOPMed reference panel (50).

### HLA class I and class II Imputation and association

HLA imputation was performed using the SNP2HLA tool (14) with genotype data from chromosome 6. A total of 25,965 SNPs were available in the ill1/CTR1 dataset, and 18,406 SNPs in the ill2/CTR2 dataset. The imputation process, based on the Type 1 Diabetes Genetics Consortium (T1DGC) reference panel (51), resulted in the imputation of 125 HLA alleles at the 2-digit level.

Association analysis between the imputed HLA variants and Elite control status was conducted using PLINK (52), employing logistic regression to assess the statistical significance of associations. A meta-analysis of the ILL1 vs CTR1 and ILL2 vs CTR2 comparisons was then performed using GWAMA (53) to combine the p-values. The threshold for statistical significance was set at 0.005 for the HLA class I or class II gene alleles.

### Stratification

To assess patterns of population structure in the ill1/CTR1 and ill2/CTR2 datasets, a principal component analysis (PCA) was conducted. The analysis aimed to quantify the genetic ancestry of the participants. A set of 510,420 informative SNPs for ill1/CTR1 and 523,850 informative SNPs for ill2/CTR2 was selected for ancestral origin determination. To mitigate the influence of linkage disequilibrium, pruning was applied using an r2 threshold of 0.3, a sliding window size of 50, and a step size of 5. Regions on chromosomes 6, 8, and 17 with high linkage disequilibrium were excluded from the analysis.

The results of the PCA confirmed the homogeneity of both the ill1/CTR1 and ill2/CTR2 datasets, consistent with the findings reported in the first ICGH publication (44).

### Association Testing by meta-analysis

For the ill1/CTR1 and ill2/CTR2 datasets, a logistic regression analysis comparing Elite Controllers (ECs) and controls (Ill1 vs CTR1 and ill2 vs CTR2) was conducted using SNPtest (54). The analysis focused on 35,552 variant dosages from MHC region under an additive model. To account for population structure and minimize potential confounding effects, the first five principal components (PCs) were included as covariates in both analyses.

Following the individual dataset analyses, a meta-analysis was performed using GWAMA software (53) to combine the p-values from each dataset. The meta-analysis aimed to identify significant associations that satisfy the following conditions: a combined p-value less than or equal to 5.10^-8^, both individual p-values less than 0.05, an infotest value greater than 0.75 (indicating a good imputation quality), and the effect sizes (OR) in the same direction across datasets. After conducting the meta-analysis on the set of 35,552 SNPs imputed in the MHC region, we identified 2,626 significant SNPs.

### SNP Stepwise regression analysis

As previously mentioned, after conducting a meta-analysis on the set of 35,552 SNPs imputed in the MHC region from the ill1/CTR1 and ill2/CTR2 datasets, we identified 2,626 significant SNPs. These SNPs were then reanalyzed step by step. At each stage, additive model regressions for ill1 vs. CTR1 and ill2 vs. CTR2 were performed using PLINK (52), incorporating the first five principal components of the PCA to account for population stratification. The best signal identified at each step was added as a covariate in the subsequent analysis. This process was repeated, with a meta-analysis using GWAMA (53) conducted after each iteration, where the next best signal was included as a covariate for the following analysis. This process was stopped at the first p-value below 0.05.

### Computation of links between MHC variants (SNPs and HLA class I and class II alleles)

To explore the relationships between SNP and HLA alleles, we computed linkage LD and R² values using PLINK (52). These calculations were applied to both SNP-SNP and SNP-HLA allele pairs. Additionally, we estimated the percentage of individuals carrying combinations of specific alleles, which is more precise than the famous D’ LD coefficient (because one knows which allele is in LD with another allele, which is not indicated by the D’ coefficient between SNPs). This percentage was determined by comparing the frequencies of two SNPs: when SNP1 was more frequent than SNP2, we computed the proportion of individuals carrying the minor allele of SNP2 also carrying the minor allele of SNP1. This percentage-based approach provides a practical measure of overlap between carriers of specific alleles, offering insight into how frequently the minor allele of a SNP is present within the carriers of a less frequent SNP minor allele, and this allows to build-up haploblock relationships. In this study, the percentage threshold for SNP2 minor allele to be part of the SNP1 minor allele haploblock is 75%.

### Covariate analyses

Covariate analyses were conducted to further investigate SNP associations. We performed them in three successive steps during this study, and the results are partially summarized in Table 1 and Supplementary Table 3:

Step 1: The 17 SNPs identified through stepwise regression were found to be linked by D’. Among these, 5 SNPs were identified for their low MAF and independence (leaves of the tree highlighted in Blue in Figure 2B), and they were part of the broader set of linked SNPs. Based on these linkage relationships, we selected these 5 SNPs or a subset of them to be used as covariates. In practice we added one by one the SNPs starting first with the ones showing LD link (Figure 2A) and stopping if the p-value became unsignificant. We then tested whether a SNP under investigation remained significant after adjusting for all these covariates.

Step 2: This analysis examined the relationships between HLA alleles. We tested whether an HLA allele remained significant after adjusting for a linked allele (identified thanks to Supplementary Table 2) as a covariate. This allowed us to see if an HLA allele was still effective for the association with EC after accounting for the effect of any linked alleles. We also performed the reciprocal test to see if the variant used as a covariate remained significant when adjusted for the original HLA allele.

Step 3: Here, we examined the relationships between the SNP minor alleles selected after Step 1 and HLA alleles (Supplementary Table 4). These associations were used to create covariates by adjusting for SNPs and HLA alleles. This allowed us to evaluate whether the tested SNP or HLA alleles remained significant when accounting for these interactions.

### Haploblock identification

To identify haploblocks linked to a specific variant (SNP or HLA allele), we used the same methodology as in our previous work (12). In brief, we performed the following steps using the CTR population: starting from a SNP or an HLA allele, we computed r ^2^ and D′ coefficients, as well as the carrier percentage (% carrier) for all SNPs within the MHC, testing a total of 35,552 SNPs. We kept haploblocks that had r^2^>0.1, D′>0.7, and % carrier > 75% with the studied allele.

### Score per individual and score per SNP allele, in each subgroup

We also analyzed the number of selected SNP minor alleles and HLA alleles for each individual using a dominant model. Each subgroup was represented by a boxplot, with each dot showing an individual’s score (Figure 3). We compared scores across different groups: CTR, EC, CV1, CV2, CV3, and between carriers and non-carriers of HLA-B*57:01. For each individual, we calculated a negative score (based on 2 negative variants) and a positive score (based on 6 positive variants).

### Biological exploration of the SNPs with genetic annotation (Annovar)

We annotated the SNPs from the haploblocks using Annovar (18) on dbSNP (avSNP150) (55) and the RefGene gene database (56). In order to include all mutants of interest, we considered in our analysis all SNPs with a r^2^ LD value greater than 0.8 with any SNP of the haploblock during the annotation process. We used SIFT (20) and Polyphen 2 (21) to predict the potential effect of amino acid substitutions on the structure and function of the protein for nonsynonymous SNPs identified.

### Transcriptional impact of the SNPs with GTEx

We used Ldexpress (57) from Ldlink (58), based on Genotype-Tissue Expression project data (GTEx, v8 release) (19), to investigate the genes significantly and differentially expressed in whole blood according to the alleles of the haploblock SNPs (as well as SNPs with r^2^ > 0.8). GTEx uses the genetic LD found in specified populations (either of European descent or of African descent) to select the SNPs of interest (with an r^2^>= 0.8 in our case), and then computes the differential transcription of genes for these SNPs among all samples (735 samples of European descent).

For each SNP, we picked up the 3 genes most differentially expressed, i.e. that exhibited the lowest p-values

## Consent to Publication

All authors have reviewed and approved the manuscript and consent to its publication.

## Author contribution

Study design: JFZ; Data curation: MR, MT, TL, JLS; Data analysis: MR, SLC, JLS, MT, RMS, RT, JFD, JFZ; Results interpretation : MR, JFD, RT, JFZ; Writing: MR, AB, SLC, RT, JFZ; Editing: All authors

## Data Availability statement

The summary statistics of the significant variants supporting the findings of this study have been deposited at the following link: https://figshare.com/s/7c63ce5803174b26f301?file=43263144.

## Ethical Approval and consent to participate

The studies involving humans were approved by Ethical approval for this study involving human genetic data was obtained for all cohorts by the local institutional review board of each group/center/study participating to the ICGH collective effort. All subjects provided written informed consent. The list of the ICGH cohorts/participants is as follows : 1. The AIDS clinical Trial Group (ACTG) in the USA 2. The AIDS Linked to the IntraVenous Experience (ALIVE) Cohort in Baltimore, USA 3. The Amsterdam Cohort Studies on HIV infection and AIDS (ACS) in the Netherlands 4. The ANRS CO18 in France 5. The ANRS PRIMO Cohort in France 6. The Center for HIV/AIDS Vaccine Immunology (CHAVI) in the USA 7. The Danish HIV Cohort Study in Denmark 8. The Genetic and Immunological Studies of European and African HIV-1+ Long Term Non-Progressors (GISHEAL) Study, in France and Italy 9. The GRIV Cohort in France 10. The Hemophilia Growth and Development Study (HGDS) in the USA 11. The Hospital Clinic-IDIBAPS Acute/ Recent HIV-1 Infection cohort in Barcelona, Spain 12. The Icona Foundation Study in Italy 13. The International HIV Controllers Study in Boston, USA 14. The IrsiCaixa Foundation Acute/Recent HIV-1 Infection cohort in Barcelona, Spain 15. The Modena Cohort in Modena, Italy 16. The Multicenter AIDS Cohort Study (MACS), in Baltimore, Chicago, Pittsburgh and Los Angeles, USA 17. The Multicenter Hemophilia Cohort Studies (MHCS) 18. The NCI Laboratory of Genomic Diversity in Frederick, USA 19. The Pumwani Sex Workers Cohort in Nairobi, Kenya, and Winnipeg, Canada 20. The San Francisco City Clinic Cohort (SFCCC) in San Francisco, USA 21. The Sanger RCC Study in Oxford, UK, and in Uganda 22. The Swiss HIV Cohort Study (SHCS), in Switzerland 23. The US military HIV Natural History Study (NHS) 24. The Wellcome Trust Case Control Consortium (WTCCC3) study of the genetics of host control of HIV-1 infection in the Gambia 25. The West Australian HIV cohort Study. The studies were conducted in accordance with the local legislation and institutional requirements.. The studies were conducted in accordance with the local legislation and institutional requirements. Written informed consent for participation was not required from the participants or the participants’ legal guardians/next of kin in accordance with the national legislation and institutional requirements.

## Acknowledgments

The authors are grateful to all the patients and medical staff who contributed to the collection of the cohorts. The authors thank Stuart Z. Shapiro (Program Officer, Division of AIDS, National Institute of Allergy and Infectious Diseases) and Stacy Carrington-Lawrence (Chair of Etiology and Pathogenesis, NIH Office of AIDS Research) who accompanied the creation of the International Collaboration on the Genomics of HIV (ICGH). The authors are also grateful to all the scientists who participated to the ICGH international effort and helped constitute this exceptional collection of data. MT is recipient of a fellowship from the Foundation FundaMental. The Laboratory GBCM and the Foundation Jean Dausset-CEPH are grateful to the program Mécénat-Santé of Mutuelles AXA for funding this research.

