## Supplementary material for "Deep analysis of the Major Histocompatibility Complex associations using covariate analysis and haploblocks unravels new mechanisms for the molecular etiology of Elite Control in AIDS": Supplementary_Material.pdf

**Supplementary Figure1. Description of the SNPs associated with EC in the MHC region obtained after stepwise regression. Representation of the SNPs with their localization in chromosome 6 GRCh38) (x-axis) and their MAF (y-axis).**

Representation of the 17 SNPs significantly associated (p<5. 10-8) with elite control in the EC vs CTR stepwise regression analysis. In blue, the SNPs whose minor allele favors elite control. In red, SNPs whose minor allele prevents elite control.

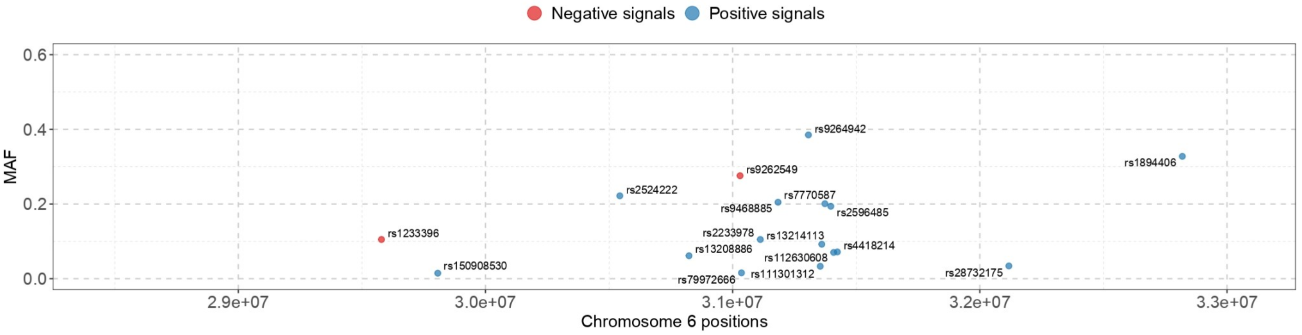


### Supplementary Tables and legends

**Supplementary Table 1. Links between the 17 SNPs obtained by stepwise regression, as computed in the CTR population.**

The SNPs are presented in decreasing maf order (from top to bottom and from left to right). The numbers correspond to the percentage of the minor allele of the SNP in the line "contained" in the minor allele of the SNP of the column (see Methods).

**Supplementary Table 2. Links between the 22 HLA alleles associated with EC (see Table2) as computed in the CTR population.**

The HLA alleles are presented in decreasing maf order (from top to bottom and from left to right). The numbers N correspond to the number of individuals carrying the HLA allele in the control population. The table provides the percentage of carriers of an HLA allele (line) who also carries the HLA allele of the column.

**Supplementary Table 3. Covariate analyses performed to eliminate interdependent SNP/HLA alleles**

**A.** Elimination by covariate analysis of the HLA alleles whose effect is dependent of the others. an allele is removed when its p-value in covariate analysis becomes non-significant. The remaining SNPs are marked in bold.

**B.** Elimination of the SNPs and HLA alleles dependent of the others. Only the HLA alleles passing the previous step A and only the SNPs remaining after the covariate analysis of Table 1 have been considered. rs28732175 explication

**Supplementary Table 4. Links between the SNP and HLA alleles (computed in CTR population)**

The table represents (A) % of HLA alleles contained in SNP minor allele as computed in the CTR group (B) % of SNP minor allele contained in the HLA alleles as computed in the CTR group.

**Supplementary Table 5. List of all the non-synonymous variants in the haploblocks of the 9 selected SNP/HLA alleles**
